# Theory for Biomolecular Catalysis in Phase-Separated Systems

**DOI:** 10.64898/2026.08.12.744453

**Authors:** Gaetano Granatelli, Samuel S. Gomez, Sudarshana Laha, Thomas C.T. Michaels, Christoph A. Weber

## Abstract

Enzymatic reactions in biomolecular condensates are often assumed to be regulated through local enrichment of reactants. However, condensates also reshape molecular transport and reaction kinetics, making it unclear how phase separation controls catalysis in living cells. Here, we develop a quantitative theory of biomolecular catalysis in phase-separated systems and find that liquid condensates can act as tunable catalytic switches, transitioning between regimes of enhanced and suppressed enzymatic activity, exhibiting optimal responses at biologically relevant condensate sizes. We show that condensate-mediated catalysis cannot be understood from reactant enrichment alone, but instead emerges from the coupled interplay of molecular partitioning, diffusive transport, and phase-dependent reaction kinetics. The strongest regulatory effects occur under rapid interphase exchange, where the spatially heterogeneous catalytic network admits a system-level Michaelis–Menten description governed by system-averaged concentrations and reaction kinetics. Our framework predicts that micron-sized condensates can either enhance or suppress enzymatic activity by up to two orders of magnitude, and that optimal catalytic regulation can emerge at condensate sizes comparable to many biomolecular condensates. These results provide experimentally testable predictions for condensate-mediated catalysis and establish quantitative principles for understanding and engineering enzyme-catalysed reactions in biomolecular condensates.

## I. INTRODUCTION

Catalysis is a cornerstone of modern chemistry and biology, in which the rate of a reaction is accelerated by a *catalyst*. Catalysts enable reactions to proceed on experimentally and biologically relevant timescales without altering thermodynamic equilibria [1]. In their absence, reaction rates are often exceedingly slow: for instance, spontaneous peptide bond cleavage would take up to four hundred years at room temperature [2], whereas hydrolysis would require up to a million years without enzymatic assistance [3]. From a theoretical perspective, catalysis reshapes the free-energy landscape of a reaction by stabilizing intermediate species or transition states [4, 5]. These principles apply broadly, from molecular to heterogeneous catalysis, such as surface-based systems [6, 7].

A particularly important class of catalysis in biological and bioengineering contexts is biomolecular catalysis [8, 9], in which macromolecules, most notably enzymes, accelerate substrate-to-product conversion [10, 11] with high molecular specificity and regulatory control [12, 13]. Such specificity is essential in complex environments, including living cells, synthetic cells, and bioreactors, where many reactions occur simultaneously [14, 15].

In these environments, spatial organization plays a central role in controlling catalytic activity. One important mechanism is phase separation, which creates distinct compartments that regulate biochemical reactions. For instance, biphasic bioreactors exploit phase separation to buffer toxic or inhibitory substrates and products, or to mediate in situ product removal [16, 17]. Similarly, cells spatially organize enzymes and other molecules using both membrane-bound domains, such as organelles, and membrane-less compartments formed by phase separation, known as *biomolecular condensates* [18–20].

Biomolecular condensates are particularly attractive due to their liquid-like nature and ability to dynamically exchange reactants across their interface with the surrounding environment [19–21]. They have been shown to regulate diverse enzyme-catalysed processes, including innate immune DNA sensing [22, 23], glycolysis [24, 25], and de novo purine biosynthesis [26], through mechanisms such as enzyme-substrate co-enrichment, reactant sequestration, and alterations of the local physicochemical environment [27–30]. Condensates have also been implicated in controlling metabolic fluxes by redirecting metabolites between competing pathways in response to cellular demands [24–27].

This raises a central question: what physicochemical mechanisms underlie condensate-mediated control of catalytic activity? A widely invoked explanation is local enrichment of reactants [31, 32]: when enzymes E and substrates S preferentially partition into a condensate, their increased local concentrations enhance the reaction rate, *r* ∝ [E][S], according to the dilute law of mass action. At first sight, reactant enrichment therefore appears sufficient to explain condensate-mediated regulation. However, this picture does not generally hold in phase-separated systems, not even locally [33, 34]. Under non-dilute conditions, reactions are governed by chemical activities *a*_*i*_ = *γ*_*i*_ · [*i*] rather than concentrations [*i*], where the activity coefficients *γ*_*i*_({[*j*]}) depend on the concentrations of all components. Most importantly, in systems at phase equilibrium, these concentration-dependent activity coefficients can offset local enrichment relative to the surrounding phase [33, 35]. Therefore, reactant enrichment and activity compensation are intrinsically coupled, such that increased local concentrations do not necessarily translate into increased catalytic activity. Consequently, condensate-mediated catalysis cannot, in general, be understood from reactant enrichment alone, despite the widespread view that condensates regulate catalysis by locally concentrating reactants.

More generally, phase separation does more than redistribute reactants in space. By altering the physicochemical environment of the catalyst, it simultaneously reshapes molecular partitioning, diffusive transport, and reaction kinetics [27, 29], such that molecular enrichment alone is insufficient to predict whether catalysis will be enhanced or suppressed. Because these effects are intrinsically coupled, the generic principles governing catalytic regulation in condensates remain elusive.

To decipher how condensates regulate catalysis, it is therefore necessary to extend the foundational frameworks of biomolecular catalysis, including Michaelis-Menten and Briggs-Haldane, to phase-separated systems. Such a theory should be quantitative and capable of addressing the catalytic effects of condensates in terms of measurable parameters accessible through currently available experimental methods, including fluorescence microscopy [36] and emerging label-free approaches [37].

In this work, we develop a theoretical framework for biomolecular catalysis in phase-separated systems. We identify the coupled interplay of measurable physico-chemical parameters, including condensate size, molecular partitioning, diffusivities, and phase-dependent reaction kinetics, as the key determinants of catalytic response. Our framework enables quantitative analysis of how condensate properties regulate catalytic readouts and predicts experimentally testable transitions between enhancement and suppression regimes, including optimal condensate sizes within the range observed in many biological systems. These results suggest that biomolecular condensates can function as tunable catalytic switches whose activity is controlled by transport, partitioning, and reaction kinetics. In the regime of rapid interphase exchange, where the strongest regulatory effects emerge, the spatially heterogeneous reaction network reduces to an effective homogeneous system described by system-averaged concentrations and reaction kinetics, yielding a Michaelis–Menten-like description of biomolecular catalysis at the system level. More broadly, our results provide experimentally testable predictions for condensate-mediated catalysis and offer a quantitative tool to decipher biochemical regulatory mechanisms based on enzymatic activity in synthetic and living cells.

## II. THEORETICAL FRAMEWORK

In this section, we extend classical Michaelis–Menten and Briggs–Haldane enzyme kinetics [38–45] to spatially heterogeneous systems with coexisting phases.

### Scaffolds and clients

To isolate the effects of phase separation on catalysis, we adopt a client-scaffold framework [33, 35], a widely used and biologically relevant description of chemical reactions occurring in biomolecular condensates [19, 46, 47]. In this framework, *scaffold* molecules control the propensity of the system to phase separate and thereby define the properties of the coexisting phases [19]. Chemically reactive species are treated as dilute *clients*, meaning that their concentrations remain small compared to those of the scaffolds [19, 48]. Consequently, clients do not significantly perturb the scaffold spatial architecture, but instead partition between coexisting phases, diffuse, and undergo chemical reactions.

Although reacting species are not always dilute in condensates, the dilute-client limit provides a minimal and experimentally relevant setting for quantifying how phase separation regulates catalysis. Within this limit, the phase-separated environment may be treated as a stationary background for client dynamics. For sufficiently sharp interfaces, client dynamics can then be described by coupled reaction–diffusion equations in each phase, supplemented by interfacial boundary conditions [35].

Situations in which reacting species significantly perturb the phase-separated scaffold background require non-dilute descriptions based on Cahn–Hilliard-type dynamics and are beyond the scope of this work [49–51].

#### Model

We consider a mixture of solvent, scaffold components, and dilute clients, where each species *i* can be described by a concentration field *c*_*i*_(**r**, *t*) that depends on space **r** and time *t*. We assume spherical symmetry such that all fields depend only on the radial position *r* = |**r**| . The mixture is considered incompressible, meaning that the molecular volumes are assumed constant.

### Scaffold phase coexistence

For clarity, we model a single scaffold species (*i* = 1) that phase-separates into two coexisting phases: a scaffold-rich condensate of radius *R* extending in 0 *< r < R* (phase I), and a surrounding scaffold-poor phase (II) occupying *R < r < L* (Fig. 1a). The framework can be easily generalized to multiple scaffolds and phases. Scaffold and solvent are treated as chemically inert and do not participate in the client reaction network. When the interfacial width of the condensate is thin compared to the bulk phases, the scaffold concentration profile *c*_1_(**r**) is stationary and well approximated by a step function, with homogeneous equilibrium concentrations 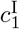 for *r < R* and 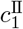 for *r > R* (Fig. 1a).

**Figure 1.**
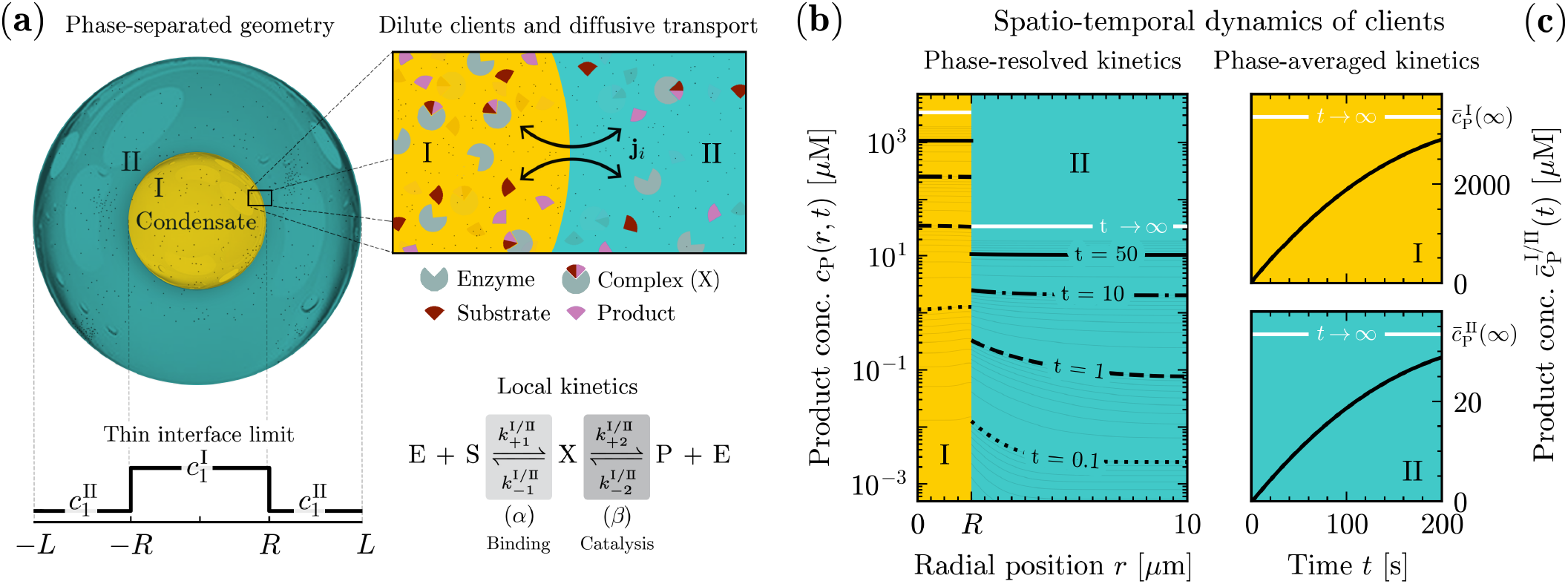
Reaction–diffusion framework for biomolecular catalysis of dilute clients in phase-separated systems. **(a) Model system.** *Geometry:* schematic of a spherically symmetric system composed of a scaffold-rich condensate (phase I) of radius *R*, coexisting with a surrounding scaffold-poor phase (phase II). The interface is assumed to be thin compared to the bulk phases (thin interface limit, bottom). *Dilute clients and transport:* dilute client species (enzyme E, substrate S, intermediate complex X, and product P) partition between the phases and are exchanged across the interface via diffusive fluxes **j**_*i*_, which satisfy flux continuity and local phase equilibrium. *Local kinetics:* within each phase, clients undergo reversible Michaelis–Menten catalysis. **(b) Phase-resolved spatio-temporal kinetics**. Representative radial concentration profiles of the product *c*_P_(*r, t*) at different times, illustrating the emergence of spatial heterogeneity during the transient dynamics. Profiles are obtained by solving the coupled reaction–diffusion dynamics governing client transport and local catalysis in the two phases (Eq. (1)). Discontinuities at the interface reflect phase-dependent partitioning, while gradients within each phase arise from the competition between local catalytic reactions and diffusive transport. At long times, diffusion relaxes spatial gradients toward the equilibrium state (white curves), determined by chemical equilibrium, phase partitioning, and global conservation constraints. **(c) Phase-averaged kinetics**. Temporal evolution of the phase-averaged product concentration 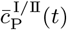 in the condensate (top) and surrounding phase (bottom). These observables are obtained by averaging the spatially resolved dynamics shown in panel (b) over each phase volume. Parameters are given in Table I.

**Table I.** Reference parameter values used throughout the manuscript unless stated otherwise.

| Symbol | Description | Value | Units |
| --- | --- | --- | --- |
| **Scaffold geometry** |  |  |  |
| $R$ | Condensate radius | 2 | $\mu\text{m}$ |
| $L$ | System radius | 10 | $\mu\text{m}$ |
| **Partitioning** |  |  |  |
| $P_E$ | Enzyme partition coefficient | 100<sup>†</sup> | – |
| $P_S$ | Substrate partition coefficient | 100 | – |
| $P_X$ | Complex partition coefficient | 100 | – |
| $P_P$ | Product partition coefficient | 100 | – |
| $\Lambda_\alpha$ | α-step kinetic partition coefficient | 10 | – |
| $\Lambda_\beta$ | β-step kinetic partition coefficient | 3000 | – |
| **Reaction kinetics<sup>‡</sup>** |  |  |  |
| $\psi_E$ | Total enzyme concentration | 0.6 | $\mu\text{M}$ |
| $\psi_S$ | Total substrate concentration | 60 | $\mu\text{M}$ |
| $k_{+1}^{\text{II}}$ | Binding rate constant | $5 \times 10^5$ | $\text{M}^{-1}\text{s}^{-1}$ |
| $k_{-1}^{\text{II}}$ | Unbinding rate constant | 5 | $\text{s}^{-1}$ |
| $k_{+2}^{\text{II}}$ | Catalytic rate constant | 0.16 | $\text{s}^{-1}$ |
| $k_{-2}^{\text{II}}$ | Reverse catalytic rate constant | $5 \times 10^{-2}$ | $\text{M}^{-1}\text{s}^{-1}$ |
| **Transport** |  |  |  |
| $D^{\text{I}}$ | Diffusivity in condensate | 4.8 | $\mu\text{m}^2\text{s}^{-1}$ |
| $D^{\text{II}}$ | Diffusivity in dilute phase | 48 | $\mu\text{m}^2\text{s}^{-1}$ |

<sup>†</sup> For the enzyme-substrate segregation regime (Fig. 6), $P_E = 0.4$ .
<sup>‡</sup> Reactant-stationary parameter $\varepsilon \equiv \psi_E/(K_S + \psi_S(0)) \approx 8 \times 10^{-3} \ll 1$ ; reversibility parameter $\delta \equiv k_{-2}\varepsilon/[k_{+1}(K_S + \psi_S(0))] \approx 1 \times 10^{-5} \ll 1$ (see Sec. II B of Supplementary Information).

### Client reaction–diffusion dynamics

The spatio-temporal evolution of dilute clients is governed by coupled partial differential equations representing reaction– diffusion processes. In particular, client concentrations obey Fickian diffusion coupled to chemical reactions [35]:

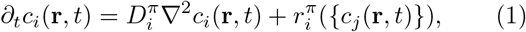

where *i, j* ≠ 1 label the client species, and *π* = I, II the phases. The first term describes Fickian transport, with phase-dependent constant diffusivities 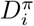 reflecting differences in transport properties between the phases. The second term describes chemical reactions through the net reaction flux 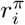 of species *i*, whose explicit form is determined by the reaction network introduced below.

### Client equilibrium partitioning

Before introducing the reaction network, we consider the diffusion-only limit 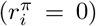, in which client concentrations relax to equilibrium values within each phase, with 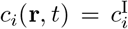 for *r < R* and 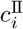 for *r > R*. The resulting equilibrium distribution of client species between the coexisting phases provides the thermodynamic baseline upon which reaction kinetics act. The relative abundances across phases are then quantified by the partition coefficient

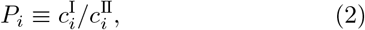

which characterizes the enrichment (*P*_*i*_ *>* 1) or depletion (*P*_*i*_ *<* 1) of species *i* in the scaffold-rich condensate. For dilute clients and fixed scaffold concentrations, *P*_*i*_ are dimensionless constants set by the energetic interaction of clients with scaffold and solvent, as well as by the scaffold concentration difference between phases [35].

### Minimal reaction network

To specify the reaction fluxes 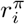, we consider a reversible single-substrate, single-product catalytic reaction occurring locally within each phase. Within the canonical framework of biomolecular catalysis [52, 53], substrate-product interconversion proceeds via two reaction steps, *η* = *α, β*, according to the reversible Michaelis–Menten mechanism

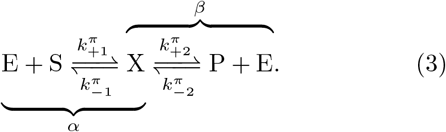

Catalysis therefore proceeds through reversible capture of substrate S by enzyme E into an intermediate complex X (*α*), followed by reversible release of enzyme E and product P from the same complex (*β*). All species are treated as dilute clients and thus evolve according to Eq. (1). These two steps are characterized by phase-dependent rate constants 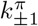 and 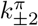, reflecting the distinct physicochemical environments of the phases. Throughout this work, the scaffold-poor values 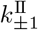 and 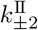 serve as reference parameters (values in Table I), while the corresponding condensate values 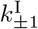 and 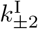 are determined through the partitioning relations introduced below.

The reversible Michaelis–Menten mechanism provides a minimal network capable of describing substrate capture, catalytic conversion, and product release [54–57]. Although we focus on this minimal model throughout the present work, the framework can be straightforwardly extended to more complex catalytic networks, including product inhibition, feedback/feed-forward regulation, and multi-substrate kinetics [40].

### Reaction fluxes and detailed balance

To ensure that the reaction kinetics is consistent with equilibrium thermodynamics, we relate the reaction fluxes to the chemical potentials of the reacting species. This guarantees that reactions relax toward thermodynamic equilibrium in the absence of external driving. The reaction fluxes 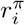 quantify the net production or consumption of each client due to reaction steps *α* and *β*. They are expressed as

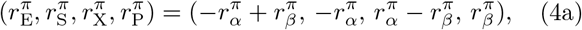

where each flux 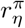 can be decomposed into a forward (+) and backward (−) contribution, 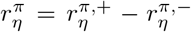. Relaxation toward thermodynamic equilibrium enforces local detailed balance in each phase [33, 58–60],

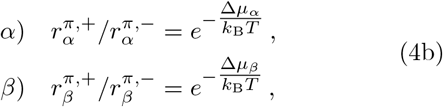

where the corresponding thermodynamic driving forces are Δ*µ*_*α*_ ≡ *µ*_X_ − (*µ*_E_ + *µ*_S_) and Δ*µ*_*β*_ ≡ (*µ*_P_ + *µ*_E_) − *µ*_X_. At chemical equilibrium, Δ*µ*_*α*_ = Δ*µ*_*β*_ = 0, and forward and backward reaction fluxes balance, 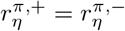.

For a client *i*, the chemical potential *µ*_*i*_ = *δF/δc*_*i*_ is defined as the functional derivative of the system free energy *F*, and takes the standard form 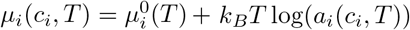, where 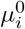 is the reference chemical potential and *a*_*i*_ the chemical activity. For dilute clients, the activity can be written as *a*_*i*_(*c*_*i*_, *T* ) = *γ*_*i*_(*c*_1_, *T* )*c*_*i*_. Notably, because the scaffold concentration *c*_1_ is uniform within each phase, the activity coefficients reduce to phase-dependent constants, 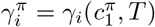 [35].

Among the possible parametrizations of forward and backward fluxes compatible with local detailed balance (Eq. (4b)), we adopt the following form:

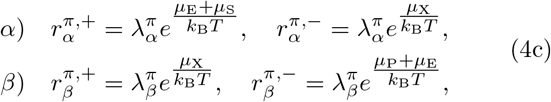

where forward and backward fluxes depend on the sums of the chemical potentials of the species participating in each reaction direction. The kinetic prefactors 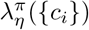 characterize the microscopic kinetics of each reaction step, and may generally depend on local composition [34]. In the dilute-client limit, however, the local environment is dominated by scaffold and solvent. Because the scaffold concentration is uniform within each phase, the kinetic prefactors then reduce to phase-dependent constants, 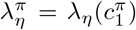, analogous to the phase-dependent activity coefficients 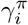 introduced above.

### Kinetic partition coefficients

The phase-dependent kinetic prefactors 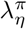 therefore reflect how the scaffold-rich and scaffold-poor environments affect the microscopic kinetics of each reaction step. In direct analogy with the partition coefficients *P*_*i*_ (Eq. (2)), we therefore define

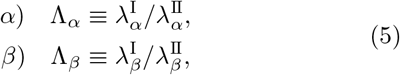

as the kinetic partition coefficients Λ_*η*_. Values Λ_*η*_ *>* 1 indicate enhanced kinetics of step *η* within the condensate, whereas Λ_*η*_ *<* 1 indicate suppression. While the equilibrium partition coefficients *P*_*i*_ quantify the redistribution of clients between phases, the kinetic partition coefficients Λ_*η*_ quantify the modulation of the microscopic kinetics of each step by the phase-separated environment.

### Effective mass-action kinetics

The thermodynamically consistent fluxes in Eq. (4c) are formulated using chemical potentials *µ*_*i*_ and kinetic prefactors *λ*_*η*_. Under the dilute-client approximation, for which the kinetic prefactors are phase-dependent constants, these fluxes can be recast in terms of concentrations *c*_*i*_, yielding an equivalent mass-action form with effective rate constants 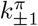 and 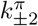 . These quantities appear directly in the reaction scheme of Eq. (3) and correspond to the kinetic parameters typically measured experimentally. Using the relation between chemical potentials and activities *a*_*i*_, the reaction fluxes can be rewritten as

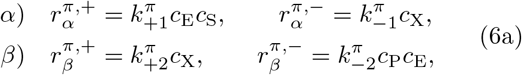

where

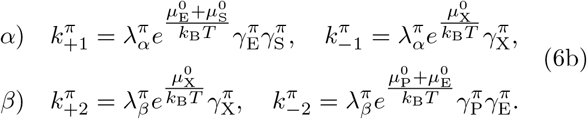

Although mathematically identical to the dilute massaction law, the quantities 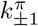 and 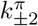 define scaffold-dependent *effective* rate constants. They combine microscopic reaction kinetics, encoded in the kinetic prefactors 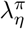, with thermodynamic contributions arising from the local scaffold environment, captured by the reference chemical potentials 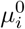 and the activity coefficients 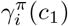 governing equilibrium partitioning (see Eq. (7b)).

The specific functional form of the effective rate constants in Eq. (6b) follows from the parametrization of the fluxes adopted in Eq. (4c). Alternative choices, such as the fully symmetric transition-state-theory parametrization, can lead to concentration-dependent effective rate constants (see Sec. I of Supplementary Information).

Evaluating Eq. (6b) in the two phases relates the effective rate constants inside the scaffold-rich condensate to those in the surrounding scaffold-poor phase:

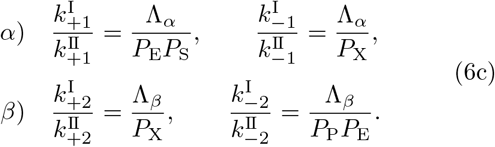

Once the effective rate constants in the scaffold-poor phase II are specified, Eq. (6c) uniquely determines the corresponding rate constants in the scaffold-rich condensate. The effects of phase coexistence are therefore fully encoded through equilibrium concentration partitioning (*P*_*i*_) and reaction kinetics partitioning (Λ_*α*_, Λ_*β*_).

### Boundary conditions

To fully specify Eq. (1), appropriate boundary conditions are required, as the presence of a sharp interface couples reaction–diffusion events across coexisting phases. The boundary conditions read:

1. *Diffusive flux continuity*: Conservation of molecule number requires radial diffusive fluxes to be continuous across the interface,

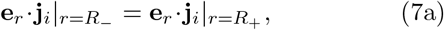

where 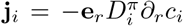 is the radial diffusive flux density, and **e**_*r*_ is the radial unit vector.
2. *Local phase equilibrium*: We assume that interfacial relaxation and transfer across the phase boundary occur on timescales much shorter than bulk diffusion and chemical reactions, such that local phase equilibrium is maintained at the interface throughout the dynamics. Accordingly, the equilibrium partition coefficient introduced in Eq. (2) applies locally at the interface, and client concentrations on either side satisfy

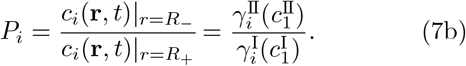
3. *System boundary* : The system boundary is impermeable, implying that diffusive fluxes vanish:

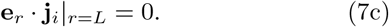
4. *Condensate centre*: Spherical symmetry requires that the diffusive fluxes at *r* = 0 vanish:

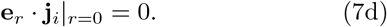

### Conclusions

In summary, phase coexistence establishes distinct physicochemical environments that regulate biomolecular catalysis through coupled effects on the concentration partitioning and kinetic partitioning of reacting client species. On one hand, scaffold phase separation redistributes clients across phases according to the equilibrium partition coefficients *P*_*i*_. On the other hand, it modifies the microscopic kinetics of individual reaction steps, as captured by the kinetic partition coefficients Λ_*α*_ and Λ_*β*_. Together, these quantities define the thermodynamic and kinetic conditions governing biomolecular catalysis of dilute clients. Although local phase equilibrium is maintained at the interface, chemical reactions generally drive the bulk concentrations away from the corresponding partitioned state. The resulting diffusive fluxes across the condensate interface alter local composition and, in turn, feed back on reaction kinetics. This mutual coupling between diffusion, partitioning, and chemical reactions is a defining feature of biomolecular catalysis in phase-separated systems and can profoundly reshape catalytic activity.

## III. PHASE SEPARATION REGULATES BIOCATALYSIS OF DILUTE CLIENTS

We now apply the framework developed above to determine how phase separation regulates biomolecular catalysis through concentration partitioning, kinetic partitioning, diffusive transport, and condensate size.

### a. Reference system without condensates

We compare catalysis in systems with and without condensates, using as reference the homogeneous limit obtained when the scaffold-rich condensate vanishes, V^I^ = 0 [35]. The relevant control parameter is the total scaffold concentration in the system, 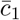, which determines whether phase coexistence occurs and, when it does, sets the condensate size. When 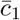 is below the equilibrium concentration of the scaffold-poor phase 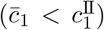, the mixture lies outside the coexistence region and remains homogeneous. In this regime, the system reduces to the scaffold-poor phase II, so that all phase-dependent quantities (concentrations, thermodynamic variables, and kinetic parameters) become spatially uniform and coincide with their scaffold-poor values. This defines the homogeneous reference system used as a benchmark throughout the phase-separated analysis. Importantly, it corresponds to the continuous limit of the phase-separated system as the condensate volume vanishes, V^I^ → 0.

For 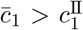, the system enters the coexistence region and phase separation occurs, giving rise to a scaffold-rich condensate of concentration 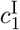 and volume

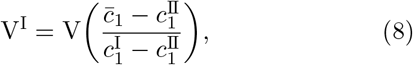

in a spherical system of volume 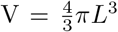 (Fig. 1(a)). The surrounding scaffold-poor phase retains concentration 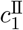 and occupies the remaining volume V^II^ = V − V^I^. Varying 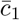 thus regulates the condensate size, while the coexisting equilibrium concentrations 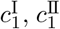 remain fixed.

### b. Setup: parameters and initial conditions

Unless stated otherwise, the results presented below are obtained by numerical solution of Eq. (1) using experimentally motivated parameter values (see Table I). To facilitate comparison with experiments, all quantities are reported in physical units, including length scales (*µ*m), concentrations (*µ*M), and time (s, min).

#### Concentration and kinetic partitioning

As a representative baseline scenario, we consider a condensate that enriches all client species relative to the surrounding dilute phase (*P*_*i*_ *>* 1) and enhances both reaction steps, with the catalytic step *β* being enhanced more strongly than the binding step *α* (Λ_*β*_ *>* Λ_*α*_ *>* 1).

#### Effective rate constants

The effective rate constants 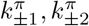 are specified by choosing representative values in the dilute phase II; the corresponding values in the condensate (I) are then uniquely determined by Eq. (6c). In particular, representative values of 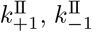, and 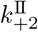 are taken from Ref. [61], which provides parameter sets consistent with experimental measurements of lysozyme and chymotrypsin [62–65]. Because experimental estimates of 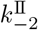 are comparatively scarce, we assign it a small but finite value such that the reverse catalytic flux remains negligible compared to the forward flux, 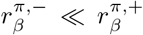. This ensures thermodynamic consistency while keeping the catalytic step effectively irreversible. All the considered rate constants lie well within the experimentally observed distributions across diverse enzymes, as reported in large-scale databases such as BRENDA [66].

#### Diffusivities

For simplicity, all client species share the same diffusivity within a given phase, *D*^*π*^. Because reductions in diffusivity within biomolecular condensates are commonly observed for macromolecules, consistent with FRAP experiments in which spatio-temporal transport models are used to extract phase-specific diffusion coefficients [67], we use a representative tenfold reduction in diffusivity inside the condensate (*D*^I^ = 10^−1^*D*^II^).

#### Initial conditions

Clients are initialized at phase equilibrium. Thus, each species *i* at *t* = 0 is spatially uniform within each phase and satisfies

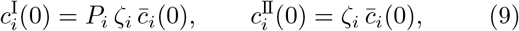

where *ζ*_*i*_(*P*_*i*_, V^I^) is the partitioning degree of species *I* which quantifies how a system-averaged concentration is distributed between coexisting phases according to the equilibrium partition coefficient *P*_*i*_ (Appendix A).

The system-averaged concentrations are defined as

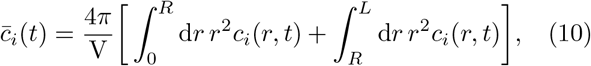

and their initial values satisfy

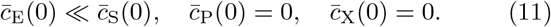

The same initial conditions are used in systems with and without condensates, ensuring that differences in catalytic response arise solely from the scaffold phase-separated environment.

#### Conserved quantities

During the ensuing dynamics, the concentrations 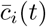 evolve in time, while

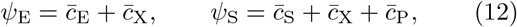

remain constant. These correspond to the conserved total enzyme and total substrate concentrations, respectively.

### 1. Spatio-temporal dynamics of clients

Having specified the model parameters and initial conditions, we now characterise the spatio-temporal evolution of client concentrations *c*_*i*_(*r, t*) governed by Eq. (1).

#### Phase-resolved kinetics

The system is initially at phase equilibrium (Eq. (9)) but out of chemical equilibrium. The onset of chemical reactions drives the system away from equilibrium, generating transient concentration heterogeneities across the system, with spatial gradients developing within each phase from the competition between local reaction kinetics and diffusive transport, as shown representatively for the product concentration in Fig. 1b. Equilibrium partitioning is continuously enforced at the condensate interface (Eq. (7b)), giving rise to discontinuities set by the partition coefficients *P*_*i*_.

During this transient regime, concentrations remain out of chemical equilibrium in the bulk. Over time, diffusion progressively relaxes these gradients, and the concentrations approach chemical equilibrium within each phase, constrained by interfacial partitioning and conservation laws. This corresponds to a steady state in which both chemical and phase equilibrium are satisfied.

#### Phase-averaged kinetics

While the client dynamics is described by spatially resolved fields *c*_*i*_(*r, t*), it is useful to introduce phase-averaged concentrations,

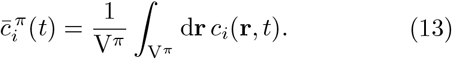

These quantities provide a coarse-grained description of the dynamics within each phase and determine, through Eq. (10), the system-averaged concentrations 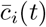. In particular, the dynamics of the phase-averaged product concentrations, 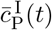 and 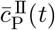, quantifies how substrate-product interconversion proceeds differently within each phase (Fig. 1c), while the system-averaged product concentration 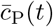 provides the experimentally relevant kinetic readout defined below: its early-time behaviour determines the initial rate, whereas its long-time relaxation towards equilibrium defines the half-time.

### A. Quantifying catalytic response: kinetic readouts

To quantify how condensates affect biomolecular catalysis relative to a condensate-free reference system, we introduce the *initial rate* and the *half-time*. These readouts probe catalytic activity at early and long times, respectively. The initial rate measures substrate-product interconversion at early times, following the establishment of a quasi–steady state, while the half-time provides a global measure of the approach to equilibrium, incorporating the cumulative effects of reversibility, enzyme saturation, and substrate depletion. For clarity, we focus on the initial rate in the main text, while long-time effects captured by the half-time are discussed in Appendix B.

### 1. Initial rate in phase-separated systems

The initial rate is defined once the system has relaxed to a quasi–steady state (QSS), characterized by an effectively stationary complex concentration and approximately constant rate of substrate-product interconversion, i.e., a nearly linear increase of product concentration with time. For the parameter values used throughout this work, the existence of a well-defined QSS regime is supported by the validity criteria for reversible Michaelis–Menten kinetics. In particular, both the reactant-stationary parameter *ε* and the reversibility parameter *δ* are much smaller than unity (see Table I), ensuring that substrate depletion and reverse product formation remain negligible during the initial transient and allowing the complex to relax rapidly to quasi–steady state (see Sec. II of the Supplementary Information).

Consistent with the definition in homogeneous systems, we define the initial rate 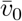 for phase-separated systems as the system-averaged rate of product formation, evaluated at the onset of the quasi–steady-state regime:

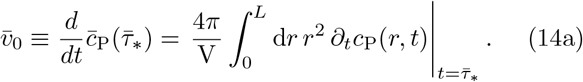

Identifying the onset of the quasi–steady-state regime is, however, non-trivial in phase-separated systems. The competition between local reaction kinetics and interphase diffusive transport generates spatial gradients within each phase (cf. Fig. 1b). As a result, the complex concentration reaches a QSS that is, in general, spatially heterogeneous across the system, so that its relaxation occurs neither uniformly in space nor synchronously in time. Unlike homogeneous systems, no single timescale characterizes the equilibration of the complex. This behaviour can be formally described by a continuum of local quasi–steady-state timescales *τ*_∗_(*r*) associated with the relaxation of the complex at each radial position *r*.

#### Coarse-grained quasi–steady state

The local quasi– steady-state timescales *τ*_∗_(*r*) are, in general, not directly accessible experimentally because measurements typically probe spatially averaged concentrations and cannot resolve fast local relaxation dynamics. We therefore introduce a *coarse-grained* quasi–steady-state timescale 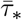 defined as the time when the system-averaged complex concentration 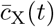 becomes stationary (Fig. 2a),

**Figure 2.**
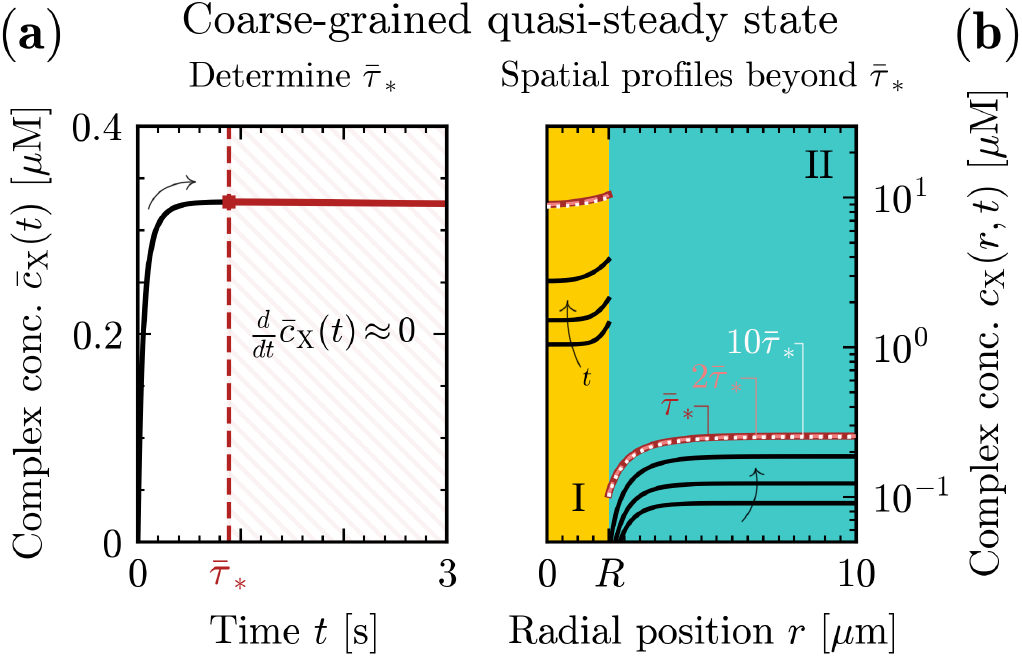
Definition of the coarse-grained quasi–steady state. **(a) Determine** 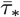. The coarse-grained QSS time is defined from the system-averaged complex concentration 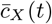 by the condition 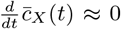. For 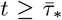, the system enters the QSS regime and 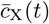 becomes effectively stationary. **(b) Spatial heterogeneity at QSS.** Representative profiles of the complex before and after 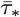. Although the concentration profile remains spatially heterogeneous, its temporal evolution is nearly stationary for 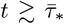, indicating that the coarse-grained criterion in (a) provides a useful proxy for the onset of local QSS dynamics. Parameter values in Table I.

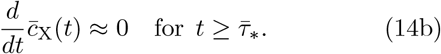

The timescale 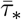 is introduced as an experimentally accessible proxy for the onset of the local quasi–steady state. Numerically, we find that the temporal evolution of the complex concentration profile becomes negligible

By 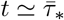, with ∂_*t*_*c*_X_(*r, t*) ≈ 0 despite the persistence of a spatially heterogeneous concentration profile (Fig. 2b).

### 2. Kinetic regimes

The initial rate in phase-separated systems depends on the interplay between chemical reactions and diffusive transport (Eq. (14)), giving rise to distinct kinetic behaviours. To characterize this interplay, we introduce reaction–diffusion length scales 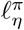 for each step *η* in each phase *π* (see Appendix C). These quantities provide order-of-magnitude measures of the distance over which concentration perturbations spread by diffusion before being significantly modified by reactions. In particular, the smallest and largest of these, *ℓ*_min_ and *ℓ*_max_ (Eq. C3), provide lower and upper bounds on the spatial extent of reaction–diffusion coupling. Whether diffusion can efficiently smooth concentration perturbations generated by reactions depends on how the reaction–diffusion length scales compare with the system geometric length scales.

Comparing the family of reaction–diffusion length scales [*ℓ*_min_, *ℓ*_max_] with the geometric length scales of the system, namely the condensate radius *R* and system size *L*, identifies three distinct kinetic regimes (Fig. 3). When reaction–diffusion and geometric length scales are comparable, diffusion and reactions compete on similar scales, generating concentration gradients throughout the system. This regime corresponds to the reaction–diffusion behaviour observed in Figs. 1b and 2b. *Reaction-limited* behaviour emerges when *ℓ*_min_ ≫ *R, L*, such that all reaction–diffusion length scales exceed the system size. Diffusion rapidly smooths concentration perturbations, maintaining equilibrium partitioning and suppressing concentration gradients within each phase [68]. This global equilibration arises from rapid molecular diffusion, whereas homogeneous enzymatic assays typically rely on stirring or convection to maintain uniform concentrations [41]. Conversely, *diffusion-limited* behaviour emerges when *ℓ*_max_ ≪ *R, L*, such that all reaction– diffusion length scales are smaller than the geometric dimensions. Concentration perturbations remain confined to thin interfacial boundary layers, making interphase transport weak and largely decoupling the dynamics of the two phases. Conditions approaching this regime may arise in highly viscous or crowded condensates, where molecular mobility is strongly reduced [67, 69].

**Figure 3.**
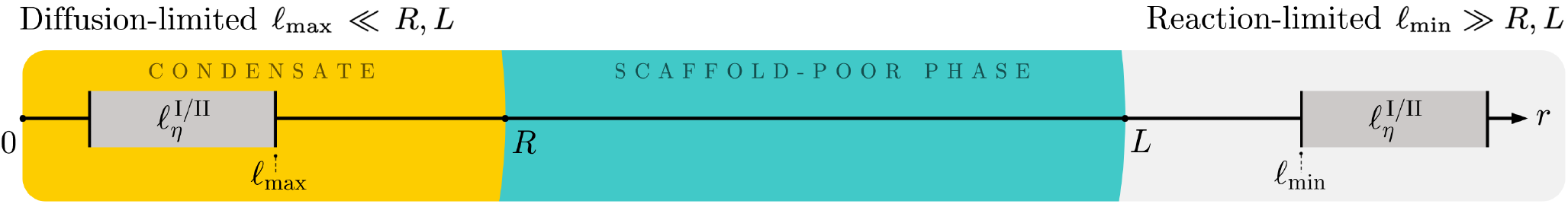
Schematic classification of kinetic regimes based on the hierarchy between reaction–diffusion and geometric length scales. The condensate radius *R* and system size *L* set the fixed geometric scales, whereas the family of reaction–diffusion length scales 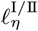 spans the interval [*ℓ*_min_, *ℓ*_max_], depicted schematically as a grey box. Two such boxes are shown to illustrate different relative positions of the entire family with respect to *R* and *L*. Reaction-limited behaviour emerges when all reaction–diffusion length scales exceed the geometric dimensions (*ℓ*_min_ ≫ *R, L*), whereas diffusion-limited behaviour emerges when all reaction–diffusion length scale remain smaller than the geometric dimensions (*ℓ*_max_ ≪ *R, L*).

The reaction-limited and diffusion-limited regimes are especially important because the underlying dynamics of the client species simplifies in the corresponding asymptotic limits, *ℓ*_min_*/R* →∞ and *ℓ*_max_*/R* → 0 respectively, enabling analytical descriptions (see Sec. III B).

### 3. Transport-controlled enhancement of the initial rate

We first examine how the initial rate varies with the initial substrate concentration across the kinetic regimes, while keeping all condensate properties fixed (Table I).

The presence of a condensate can strongly enhance the initial rate relative to the reference homogeneous system, but the magnitude of this enhancement is controlled by diffusive transport (Fig. 4a). Under diffusion-limited conditions, interphase transport is weak, so that enhanced kinetics within the condensate cannot be efficiently supplied with reactants from the surrounding phase. Consequently, condensate-enhanced kinetics remains only weakly expressed at the system level. As reaction–diffusion length scales increase (corresponding to higher diffusivities), transport limitations are progressively alleviated and the system-level enhancement becomes more pronounced. Under reaction-limited conditions, diffusion rapidly redistributes clients across phases and continuously replenishes the reactants consumed within the condensate, maintaining equilibrium partitioning and suppressing concentration gradients. This allows the condensate-enhanced kinetics to be fully expressed at the system level, resulting in a strong enhancement of the initial rate.

**Figure 4.**
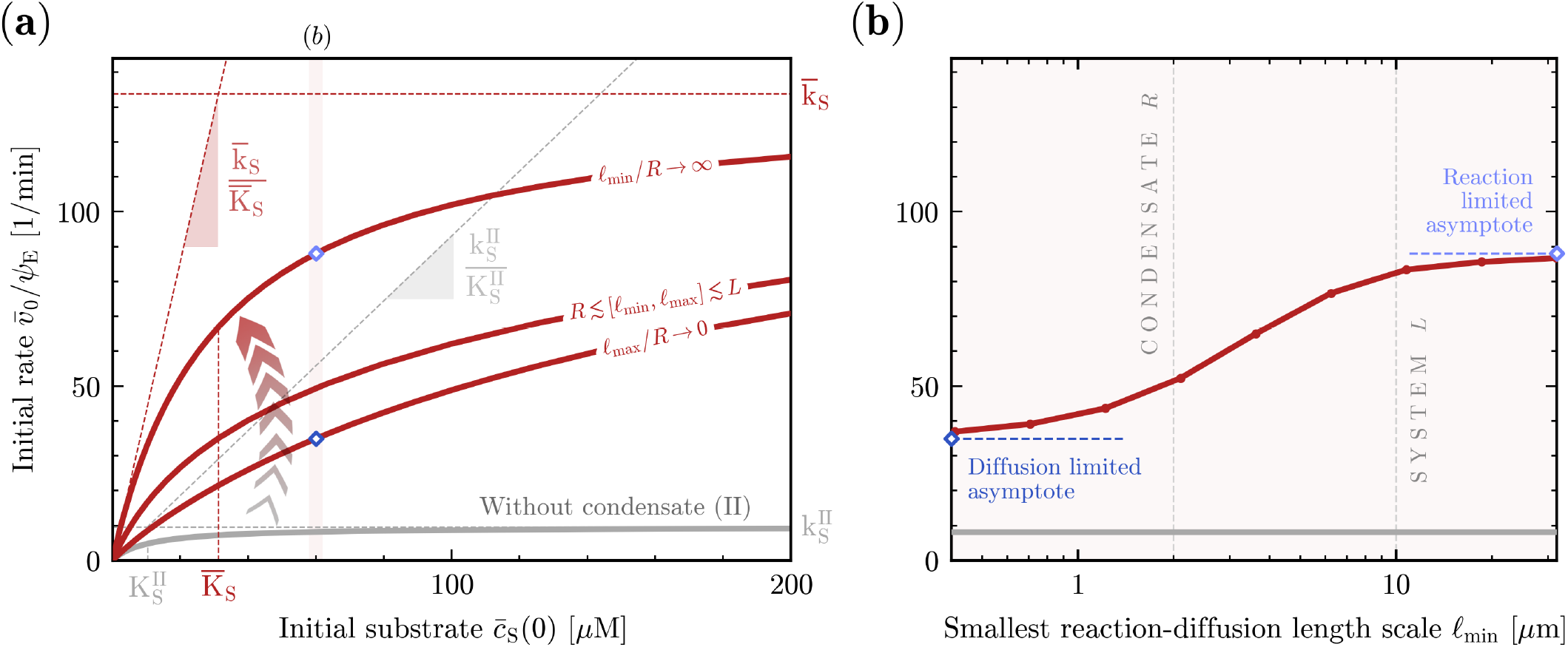
Transport-controlled enhancement of the initial rate mediated by condensates. For fixed condensate properties, a phase-separated condensate can strongly enhance the initial rate relative to the condensate-free reference system. The magnitude of this enhancement is controlled by diffusive exchange of clients between phases. We compare systems with condensates (red) to system without condensates (gray). **(a)** Enzyme-normalized initial rate 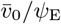 as a function of the initial substrate concentration 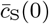. Red curves correspond to representative kinetic regimes: diffusion-limited (lower), reaction– diffusion (intermediate), and reaction-limited (upper). In particular, the diffusion-limited (lower) and reaction-limited (upper) curves are obtained directly from Eqs. (19) and (28), respectively, whereas the intermediate curve is obtained numerically from the full reaction–diffusion model (Eq. (14)). For the homogeneous reference system and the asymptotic reaction-limited system, dashed lines indicate the corresponding Michaelis–Menten parameters and low-substrate slopes, obtained from the homogeneous and coarse-grained analytical descriptions derived in the Supplementary Information and Sec. III B, respectively. The shaded vertical band indicates the substrate concentration used in panel (b). **(b)** Enzyme-normalized initial rate 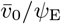 as a function of the smallest reaction–diffusion length-scale *ℓ*_min_. Increasing *ℓ*_min_ continuously shifts the system from diffusion-limited to reaction-limited behaviour. The two ends of the curve approach the corresponding diffusion-limited and reaction-limited asymptotic limits. Parameter values are given in Table I.

#### Crossover between kinetic regimes

The enhancement increases continuously as the system transitions from diffusion-limited to reaction-limited behaviour (Fig. 4b). To probe the transition, we uniformly rescale all client diffusivities while keeping the kinetic parameters fixed. Because reaction–diffusion length scales increase with diffusivity (Eq. (C2)), this shifts the entire family of reaction–diffusion length scales 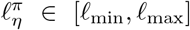 relative to the fixed geometric length scales *R* and *L*, while preserving their internal ordering. Increasing diffusivities therefore shifts the family 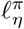 toward larger values, whereas decreasing diffusivities has the opposite effect.

As *ℓ*_min_ is decreased, progressively more reaction– diffusion length scales become smaller than *R* and *L*. This drives the system toward diffusion-limited behaviour, and in the asymptotic limit *ℓ*_max_*/R* → 0 all reaction–diffusion length scales are much smaller than the condensate radius. Conversely, as *ℓ*_min_ is increased, the entire family 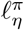 shifts to larger values. Once all reaction–diffusion length scales exceed the geometric di-mensions, reaction-limited behaviour emerges, approaching the asymptotic limit *ℓ*_min_*/R* → ∞. The initial rate therefore interpolates continuously between the diffusion-limited and reaction-limited asymptotes, demonstrating that transport controls how strongly condensate-localized kinetics is expressed at the system level.

### 4. Regulation of the initial rate by condensate properties

Having established that condensates can strongly regulate the initial rate relative to a reference homogeneous system without condensates, and that the magnitude of this regulation is controlled by diffusive transport, we now examine how it depends on condensate properties.

We vary the condensate volume fraction V^I^*/*V and the kinetic partition coefficient of the catalytic step, Λ_*β*_, while keeping the kinetic partition coefficient of the binding step fixed at Λ_*α*_ *>* 1. This isolates the contribution of catalytic turnover (reaction step *β*) from effects associated with capture and binding (reaction step *α*).

We further compare two distinct localization regimes: enzyme-substrate co-localization (*P*_E_ *>* 1, *P*_S_ *>* 1), in which both species are enriched in the condensate, and enzyme-substrate spatial segregation (*P*_E_ *<* 1, *P*_S_ *>* 1), in which the substrate is enriched while the enzyme is excluded. In both cases, all the other clients remain enriched within the condensate (*P*_S_ *>* 1, *P*_X_ *>* 1, *P*_P_ *>* 1). Together, these parameters determine whether condensates enhance or suppress the initial rate, and whether optimal condensate sizes emerge.

### a. Relative initial rate

To quantify condensatemediated regulation relative to the reference homogeneous system, we introduce the relative initial rate

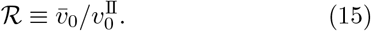

Because the initial rate depends on the kinetic regime, we focus here on reaction-limited behaviour for three reasons. First, rapid diffusion allows condensate-localized reaction kinetics to be fully expressed at the system level. Consequently, reaction-limited systems exhibit the largest deviations from the homogeneous reference system (cf. Fig. 4). Second, this regime provides a relevant reference in many biological contexts, as soluble enzymes and metabolites often equilibrate across condensates on timescales comparable to or faster than catalytic turnover [70, 71], implying that transport is effectively faster than reaction kinetics. Third, reaction-limited behaviour admits a particularly simple analytical description (Sec. III B). The phase-separated reaction network can be coarse-grained into an effective homogeneous system governed by system-averaged concentrations and rate constants. As a result, ℛ factorizes into kinetic and enzyme-binding contributions, making the effects of concentration and kinetic partitioning explicit.

### b. Enzyme-substrate co-localization

When enzyme and substrate co-localize within the condensate (*P*_E_ *>* 1, *P*_S_ *>* 1), concentration partitioning enriches both species within the condensate, while kinetic partitioning enhances the catalytic turnover of the resulting complexes. The interplay between these two effects generates a non-trivial dependence of the relative initial rate ℛ on volume fraction V^I^*/*V (Fig. 5a), leading to the emergence of optimal condensate sizes that maximize enhancement of the initial rate. In particular, we find that a critical catalytic-step kinetic partition coefficient, 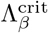, separates monotonic regulation from the emergence of optimal enhancement. The origin of this behaviour can be understood by examining the small-condensate and large-condensate limits (V^I^*/*V → 0 and V^I^*/*V → 1).

**Figure 5.**
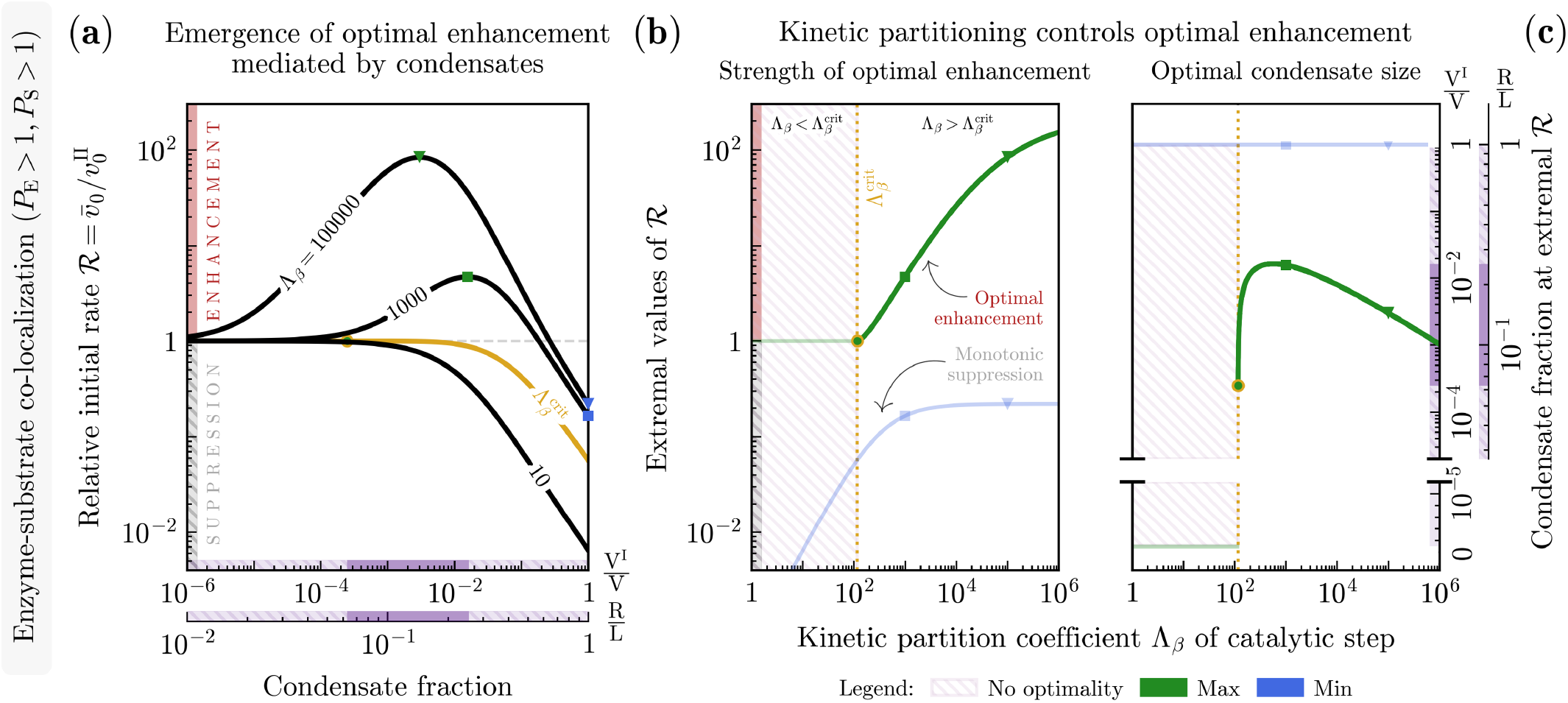
Optimal condensate-mediated enhancement of the initial rate under enzyme-substrate co-localization. Under reaction-limited kinetics and enzyme-substrate co-localization, concentration partitioning and kinetic partitioning cooperate locally but contribute differently at the system level, yielding optimal enhancement at intermediate condensate sizes. Coloured guide bars act as visual aids only: red denotes enhancement (ℛ *>* 1), grey denotes suppression (ℛ *<* 1), and purple denotes condensate fraction. Solid segments indicate parameter regions where an optimum exists (optimal regulation), whereas hatched segments indicate monotonic regulation without an optimum. **(a)** Relative initial rate ℛ as a function of both the condensate volume fraction V^I^*/*V and the corresponding radius fraction *R/L*, for different values of Λ_*β*_ (black curves). A critical catalytic-step kinetic partition coefficient 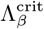 (gold curve) separates monotonic suppression from optimal enhancement. Green markers denote maxima of ℛ, whereas blue markers denote minima. In the co-localization regime, minima always occur in the large-condensate limit V^I^ → V. **(b)** Extremal values associated with the curves shown in panel (a), highlighting the strength of optimal regulation. Green and blue curves show the maximal and minimal values of ℛ, respectively. In the co-localization regime, minima coincide with the large-condensate limit. **(c)** Condensate fraction associated with the extrema shown in panel (a), reported in terms of both the volume fraction V^I^*/*V and the corresponding radius fraction *R/L*. Above 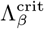, an optimal condensate size emerges and varies non-monotonically with Λ_*β*_ (green branch). Parameter values in Table I.

#### Small-condensate limit

When V^I^*/*V → 0, the condensate occupies a vanishing fraction of the total system and thus contributes negligibly to both system-averaged concentrations and system-averaged kinetics. Consequently, the phase-separated system becomes indistinguishable from a homogeneous phase-II system, and ℛ → 1.

For 0 *<* V^I^*/*V ≪ 1, enzyme and substrate become colocalized within a small condensate where concentration partitioning enriches their abundances and kinetic partitioning enhances both binding and catalysis. A critical catalytic-step kinetic partition coefficient, 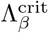, marks when kinetic partitioning becomes sufficiently strong to enhance the system-level initial rate for a given concentration partitioning, and thus separates the two possible system-level responses upon formation of a small condensate: the small-condensate slope of ℛ with respect to condensate size is negative for 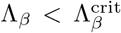 and positive for 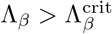 (see Sec. III B and Appendix E).

#### Large-condensate limit

As the condensate occupies an increasingly large fraction of the system volume (V^I^*/*V → 1), concentration partitioning and kinetic partitioning influence the system-level response in distinct ways. For concentration partitioning, the condensate enrichment relative to the system average, 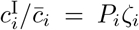, decreases from *P*_*i*_ to unity as V^I^*/*V increases from 0 to 1 (Appendix A). Consequently, concentration partitioning progressively loses its ability to enrich condensate-localized species relative to the system average. By contrast, kinetic partitioning remains explicitly encoded in the system-averaged rate constants even in the large-condensate limit (Appendix A). Thus, the concentration-partitioning advantage associated with enzyme-substrate co-localization is progressively lost, whereas kinetic partitioning continues to influence the effective reaction kinetics. This kinetic contribution alone, however, cannot compensate for the loss of concentration enrichment. As a result, irrespective of Λ_*β*_, the slope of ℛ remains negative for sufficiently large condensates (cf. Fig. 8), implying that further condensate growth always reduces the relative initial rate in this limit.

#### Optimality

The combination of these two limits explains the emergence of optimal regulation: for Λ_*β*_ *>* Λ^crit^, the slope of ℛ is positive for small condensates but negative for large ones (see Fig. 8). Consequently, ℛ develops a maximum ℛ∗ at a size V^I,∗^, defined by 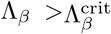 Analytical conditions for the existence of the maximum directly yield the expression for the critical value 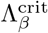 ^t^ (see Appendix E).

For 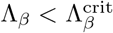, the relative initial rate remains monotonic and below unity for all condensate sizes; varying Λ_*β*_ therefore only changes the magnitude of suppression. For 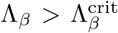, the optimal response ℛ^∗^ increases continuously with Λ_*β*_, reaching enhancements approaching two orders of magnitude relative to the homogeneous reference system (Fig. 5b). Remarkably, these strong enhancements occur for condensate radii in the micrometre range, with optimal sizes lying between *R* ≈ 0.6 and 3 *µ*m (Fig. 5c), comparable to those observed for many biomolecular condensates.

### c. Enzyme-substrate spatial segregation

When enzyme and substrate are spatially segregated (*P*_E_ *<* 1, *P*_S_ *>* 1), concentration partitioning and kinetic partitioning act in opposition. Concentration partitioning enriches substrate within the condensate but depletes enzyme, reducing the formation of reactive complexes, whereas kinetic partitioning enhances the catalytic turnover of those complexes that do form. As a result, condensate-mediated regulation displays a richer dependence on condensate size than in the co-localization regime and can give rise to monotonic suppression, optimal suppression, monotonic enhancement, and optimal enhancement (Fig. 6a). This behaviour can again be understood from the small-condensate and large-condensate limits (V^I^*/*V → 0 and V^I^*/*V → 1).

**Figure 6.**
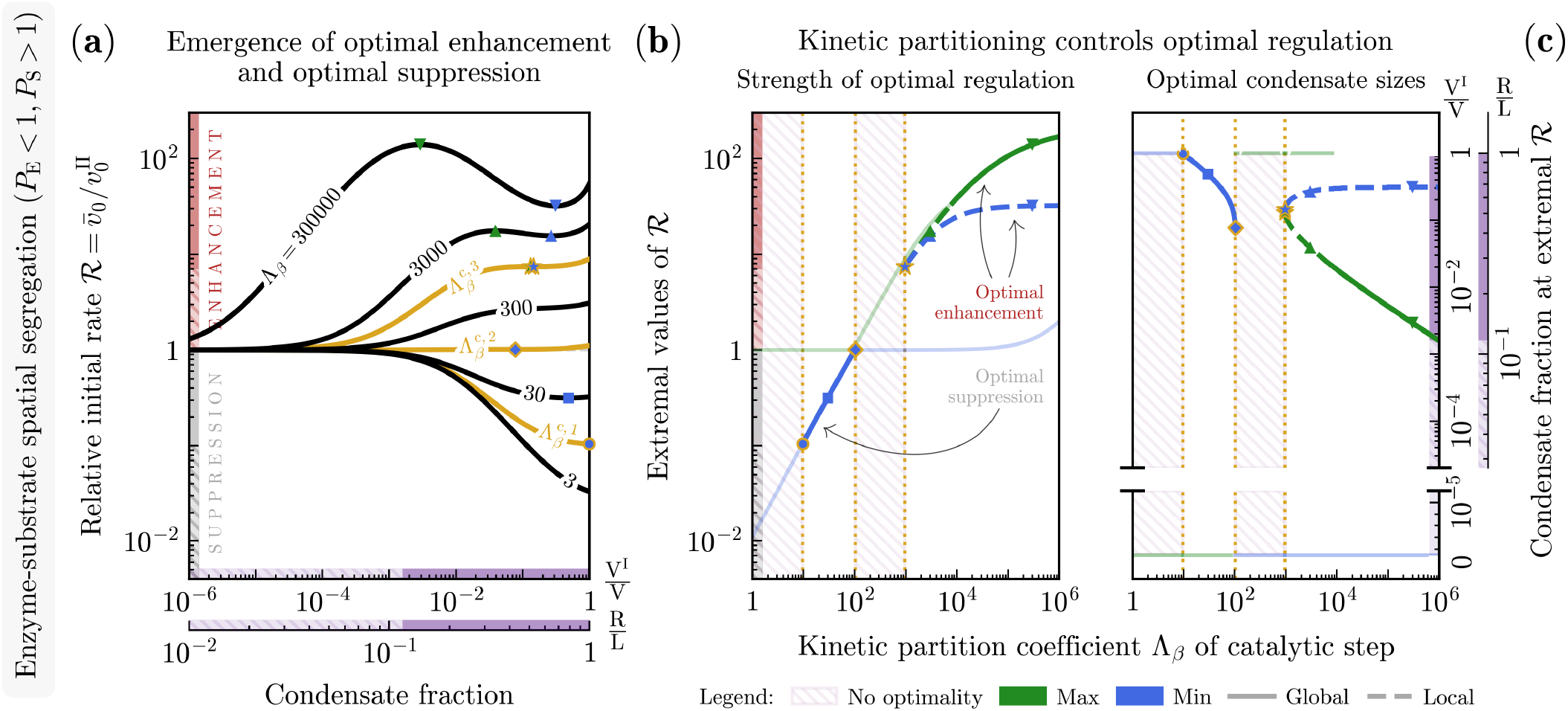
Optimal condensate-mediated enhancement and suppression of the initial rate under enzyme-substrate spatial segregation. Under reaction-limited kinetics and enzyme-substrate spatial segregation, concentration partitioning and kinetic partitioning act in opposition, generating regimes of monotonic and optimal enhancement as well as monotonic and optimal suppression. Coloured guide bars act as visual aids only: red denotes enhancement (ℛ *>* 1), grey denotes suppression (ℛ *<* 1), and purple denotes condensate fraction. Solid segments indicate parameter regions where an optimum exists (optimal regulation), whereas hatched segments indicate monotonic regulation without an optimum. **(a)** Relative initial rate ℛ as a function of both the condensate volume fraction V^I^*/*V and the corresponding radius fraction *R/L*, for different values of Λ_*β*_ (black curves). Three critical catalytic-step kinetic partition coefficients, 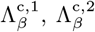, and 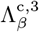 (gold curves) separate four qualitative regimes: monotonic suppression, optimal suppression, monotonic enhancement, and optimal enhancement. Green and blue markers denote maxima and minima of ℛ, respectively. **(b)** Extremal values associated with the curves shown in panel (a), highlighting the strength of optimal regulation. Green and blue curves show the maximal and minimal values of ℛ, respectively. Optimal suppression exists only for 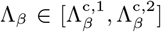, whereas above 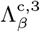 both extrema lie above unity. **(c)** Condensate fractions associated with the extremal values shown in panel (a), reported in terms of both the volume fraction V^I^*/*V and the corresponding radius fraction *R/L*. Above 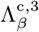, two c to the maximum and minimum of ℛ. Parameter values in Table I.

#### Small-condensate limit

When V^I^*/*V → 0, the phase-separated system becomes indistinguishable from the homogeneous phase-II reference system and ℛ → 1. For 0 *<* V^I^*/*V ≪ 1, substrate becomes enriched within the condensate whereas enzyme becomes depleted from it.

Consequently, enzyme-substrate encounters are reduced, tending to suppress the system-level initial rate. Kinetic partitioning counteracts this effect by enhancing the catalytic turnover of the complexes that do form within the condensate. The balance between these opposing effects determines whether the formation of a small condensate decreases or increases the relative initial rate; correspondingly, the small-condensate slope of ℛ with respect to condensate size can be either negative or positive.

#### Large-condensate limit

As in the co-localization regime, concentration partitioning becomes progressively less effective as V^I^*/*V → 1, whereas kinetic partitioning remains encoded in the system-averaged kinetics. Unlike the co-localization regime, however, the kinetic enhancement generated by Λ_*β*_ can completely compensate for the suppression resulting from enzyme depletion. Consequently, large condensates are not necessarily suppressive and may either suppress or enhance the relative initial rate, depending on the strength of Λ_*β*_; accordingly, the large-condensate slope of ℛ with respect to condensate size can also be either negative or positive.

#### Optimality

A key difference from the co-localization regime is that spatial segregation permits sign changes in both the small-condensate and large-condensate asymptotic slopes as Λ_*β*_ increases. By contrast, in the co-localization regime only the small-condensate slope changes sign, whereas the large-condensate slope remains negative for all Λ_*β*_. Although the asymptotic slopes are not sufficient to determine the existence of extrema in the segregation regime, where multiple extrema may occur, the ability of both asymptotic slopes to switch sign provides intuition for its richer regulation landscape compared to the co-localization regime. In particular, four qualitatively distinct forms of regulation emerge, separated by three critical values of the catalytic-step kinetic partition coefficient 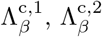, and 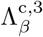 (Fig. 6a).

For 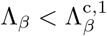, the response is monotonic and suppressive for all condensate sizes, whereas for 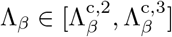 it is monotonic and enhancing. Optimal suppression arises for 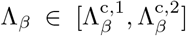, where ℛ develops a minimum below unity, while optimal enhancement emerges for 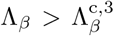, where ℛ develops both a minimum and a maximum above unity at distinct condensate sizes (Fig. 6b). For the experimentally relevant parameter values considered here (Table I), optimal suppression reaches at most about one order of magnitude relative to the homogeneous reference system, whereas optimal enhancement approaches two orders of magnitude, comparable to that observed in the co-localization regime. Thus, although enzyme depletion strongly opposes catalysis, sufficiently strong kinetic enhancement can more than compensate for this effect and generate catalytic responses far exceeding those of systems without condensates. Remarkably, the condensate radii associated with both optimal enhancement and optimal suppression lie in the micrometre range, with optimal enhancement occurring for *R* ≈ 1 − 6 *µ*m and optimal suppression for *R* ≈ 4 − 10 *µ*m (Fig. 6c), comparable to those reported for many biomolecular condensates.

The enzyme-substrate segregation regime therefore exhibits two features absent from enzyme-substrate co-localization. First, condensates can mediate both optimal suppression and optimal enhancement of the initial rate. Second, sufficiently strong catalytic enhancement can eliminate suppression altogether, allowing condensates to enhance the initial rate for all condensate sizes.

### B. Closed-form expressions for diffusion-limited and reaction-limited biocatalysis

The numerical results presented above reveal that biomolecular catalysis in phase-separated systems is strongly regulated by the interplay between diffusive transport and chemical reactions, controlling both how the initial rate changes across kinetic regimes and how it responds to variations in condensate properties.

These observations can be analytically understood by considering the asymptotic limits associated with the diffusion-limited and reaction-limited regimes introduced in Sec. III A. These regimes are identified by the hierarchy between the geometric length scales *R* and *L* and the family of reaction–diffusion length scales 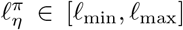. Specifically, we analyse the asymptotic limits *ℓ*_max_*/R* →0 and *ℓ*_min_*/R* → ∞, at fixed *R/L*. In both cases, the reaction–diffusion equations reduce to ordinary differential equations, yielding simplified reaction schemes and closed-form expressions for the initial rate.

### 1. Diffusion-limited kinetics

The analytical reduction presented below corresponds to the asymptotic limit *ℓ*_max_*/R* → 0.

#### a. Phase-resolved description

In this limit, the spatially resolved reaction–diffusion dynamics (Eq. (1)) reduces to a phase-resolved temporal description,

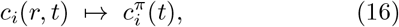

where the phase concentrations 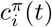 become asymptotically independent dynamical variables associated with each phase. Accordingly, the coupled partial differential equations reduce to independent ordinary differential equations within each phase,

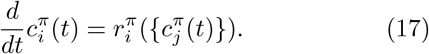

The reaction fluxes 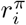 retain the same mass-action form as in Eq. (6a), evaluated for spatially homogeneous concentrations 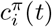 in each phase. The effective rate constants 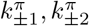 are still given by Eq. (6c). The system thus reduces to two homogeneous, kinetically isolated compartments governed by phase-specific kinetics. The homogeneous Michaelis–Menten theory derived in the Supplementary Information (Sec. II) therefore applies independently within each phase.

#### b. Initial rate

Consistent with this phase-resolved reduction, the quasi–steady-state dynamics likewise reduces to a phase-specific relaxation, such that the complex concentration in each phase relaxes on a uniform phase-specific timescale 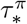. The QSS approximation thus applies independently in each phase, and the system-level initial rate (Eq. (14)) reduces to a volume-weighted sum of phase-specific contributions,

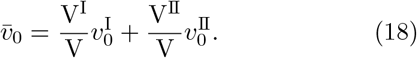

Each phase-specific initial rate, 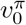, is therefore determined independently by the corresponding phase-specific kinetics and follows the same Michaelis–Menten form as in homogeneous media, with phase-specific turnover rates (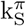 and 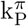), Michaelis constants (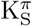 and 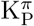), and equilibrium constant 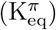. We use the factorized form of the initial rate that explicitly separates the respective roles of kinetics, thermodynamics, and enzyme binding in biomolecular catalysis [72–74],

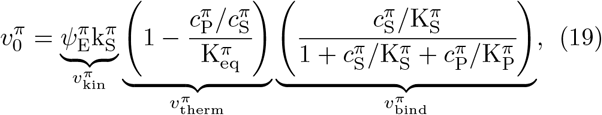

where 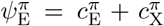 is total enzyme concentration in phase *π*. The terms 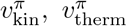, and 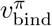 represent the phase-specific counterparts of the kinetic, thermodynamic, and enzyme-binding contributions appearing in the homogeneous reversible Michaelis–Menten theory discussed in the Supplementary Information.

Eq. (18) provides the system-averaged initial rate 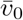 in the diffusion-limited asymptotic limit (*ℓ*_max_*/R* → 0), shown as the lower curve in Fig. 4a. The highlighted point on this curve corresponds to the initial substrate concentration used in panel (b), and its value is reproduced by the diamond marker in Fig. 4b.

### 2. Reaction-limited kinetics

The analytical reduction presented below corresponds to the asymptotic limit *ℓ*_min_*/R* → ∞.

#### a. Phase-resolved description

In this limit, as for the diffusion-limited kinetics, the spatially resolved reaction–diffusion dynamics (Eq. (1)) can first be reduced to a phase-resolved temporal description (Eq. (16)). In particular, spatial homogeneity within each phase allows the coupled partial differential equations to be integrated over the phase volumes, yielding phase-averaged balance equations [33],

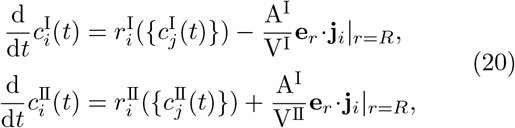

where A^I^ = 4*πR*^2^ is the condensate interfacial area. The second term represents the volume-normalized exchange of species *i* across the interface. In contrast to the diffusion-limited case (Eq. (17)), the dynamics in the two phases remain coupled through the volume-normalized interfacial transport terms appearing in Eq. (20), which continuously exchange clients between phases. The reaction fluxes 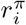 retain the same mass-action form as in Eq. (6a), evaluated for spatially homogeneous concentrations 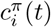 in each phase. The effective rate constants 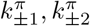 are still given by the expressions in Eq. (6c).

#### b. System-averaged description

Unlike under diffusion-limited kinetics, reaction-limited kinetics admit a further reduction beyond the phase-resolved description, allowing the dynamics to be expressed entirely in terms of system-averaged quantities. Because concentrations remain spatially homogeneous within each phase, the local phase-equilibrium condition at the condensate interface (Eq. (7b)) extends throughout the bulk of the phases. Consequently, the equilibrium partition coefficient *P*_*i*_ determines the ratio of the bulk concentrations in the two phases at all times. Moreover, the general definition of the system-averaged concentrations (Eq. (10)) reduces to

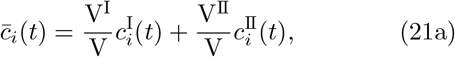

which expresses conservation of total material. Combining Eq. (21a) with the equilibrium partition coefficient (Eq. (2)) yields the phase equilibrium conditions

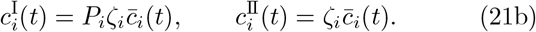

Eq. (21b) generalizes the phase equilibrium initial conditions (Eq. (9)) from *t* = 0 to the entire dynamics, thereby expressing the phase concentrations entirely in terms of the system-averaged concentrations.

##### Coarse-graining of concentrations

As a consequence, the phase concentrations 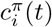 are no longer independent dynamical variables, but are fully determined by the system-averaged concentrations 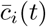. The phase-resolved description therefore admits the coarse-graining

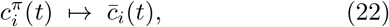

which directly follows from phase equilibrium and conservation of total material (Eq. (21)) and gives rise to the concentration partitioning degrees *ζ*_*i*_. Substituting Eq. (21b) into the phase-resolved dynamics yields

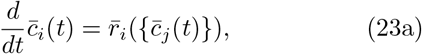

where the interfacial transport terms cancel upon summation of the phase-averaged balances owing to flux continuity at the interface (Eq. (7a)) together with conservation of molecule number. The system-averaged reaction fluxes 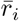 are given by the volume-weighted sum of the phase-specific contributions appearing in Eq. (20),

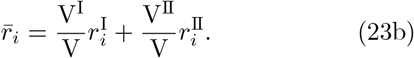

##### Coarse-graining of reaction kinetics

The reduction to system-averaged concentrations enables an analogous coarse-graining of the reaction kinetics. Rewriting the phase-resolved reaction fluxes 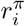 in terms of system-averaged concentrations 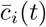,

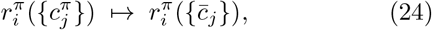

introduces phase-specific *apparent* rate constants, whose volume-weighted averaging yields the corresponding system-level kinetic parameters governing the coarse-grained description. The explicit form of the resulting system-level kinetic parameters depends on the underlying reaction mechanism. We illustrate this construction below for reversible Michaelis–Menten reaction mechanism (see Appendix D for the derivation).

Thus, under reaction-limited conditions, both concentrations and reaction kinetics admit a common system-level representation based on volume-weighted quantities. As a result, the phase-separated reaction network can be reduced to an effective homogeneous description governed entirely by system-averaged quantities. This constitutes the central result of the reaction-limited regime: rapid diffusion across the condensate interface enables a complete coarse-graining of phase-separated kinetics into a system-level framework formally analogous to the dilute mass-action description of a homogeneous reaction network. Importantly, this coarse graining procedure is not specific to the Michaelis–Menten reaction mechanism and can be applied more generally to reaction networks that admit a description in terms of time-independent phase-specific kinetic parameters.

##### Michaelis–Menten kinetics

For the Michaelis–Menten reaction mechanism, the kinetic coarse-graining introduced above gives rise to phase-specific apparent rate constants 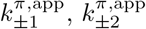, whose volume-weighted averaging yields the system-averaged rate constants 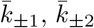:

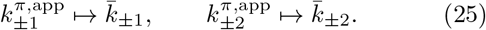

The resulting system-averaged dynamics can then be represented by the *coarse-grained reaction scheme*

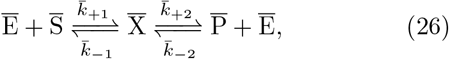

where the barred species denote system-averaged concentrations 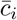 (Eq. (21a)), and the corresponding systemaveraged rate constants 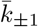 and 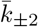 read

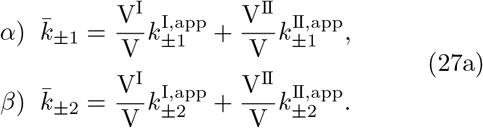

The relation between the phase-specific apparent rate constants 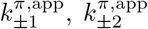 and the phase-specific effective rate constants 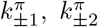 (Eq. (6c)) is discussed in Appendix D. Using the resulting expressions, the system-averaged rate constants in Eq. (27a) can be recast as renormalized versions of the corresponding values in the reference homogeneous system (i.e., *π* = II),

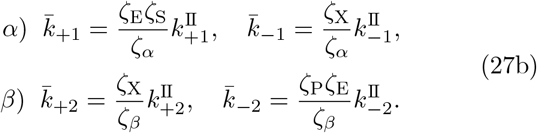

The concentration partitioning degrees *ζ*_*i*_(*P*_*i*_, V^I^) originate from phase equilibrium and conservation of total material, and quantify how the redistribution of client species between coexisting phases contributes to the system-averaged rate constants. The kinetic partitioning degrees *ζ*_*η*_(Λ_*η*_, V^I^) emerge from the reaction-limited coarse-graining of the phase-resolved kinetics, quantifying how the microscopic reaction kinetics in coexisting phases contribute to the system-averaged rate constants. Although the partitioning degrees *ζ*_*i*_ and *ζ*_*η*_ have distinct physical origins, they enter the system-averaged rate constants on an equal mathematical footing.

To conclude, biomolecular catalysis under reaction-limited kinetics retains the same Michaelis–Menten structure as a homogeneous system. All effects of phase separation enter through the concentration and kinetic partitioning degrees, *ζ*_*i*_ and *ζ*_*η*_, which renormalize the effective rate constants of the coarse-grained reaction scheme.

#### c. Initial rate

Consistent with the coarse-graining of the dynamics from the spatial level to the phase level and ultimately to the system level, the quasi–steady-state description admits an analogous reduction. Because the dynamics reduce to an effective homogeneous reaction scheme, the QSS relaxation can be characterized by a single coarse-grained timescale 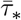, and the QSSA applies directly to the coarse-grained reaction scheme (Eq. (26)).

The initial rate admits the same system-level coarsegraining as the dynamics. Although the volume-weighted representation (Eq. (18)) remains formally valid, phase equilibrium and kinetic coarse-graining allow the initial rate to be recast directly as a closed Michaelis–Menten rate law governed by system-averaged quantities, with all effects of phase separation encoded in the renormalized turnover rates (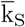 and 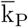), Michaelis constants (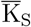 and 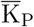), and equilibrium constant 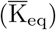. Using the same factorized form that separates kinetic, thermodynamic, and enzyme-binding contributions, the system-level initial rate 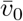 becomes

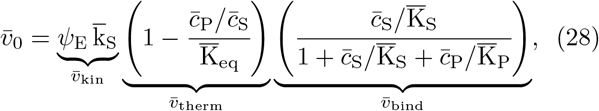

where the terms 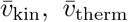, and 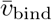 represent the coarse-grained counterparts of the corresponding kinetic, thermodynamic, and enzyme-binding contributions.

##### Kinetic contribution

For the two-step reaction mechanism considered here, the kinetic contribution 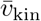 is determined by the system-averaged substrate turnover rate 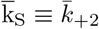, such that

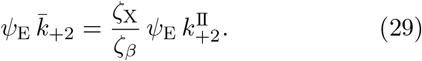

The kinetic contribution 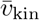 is thus obtained by renormalizing the homogeneous kinetic capacity 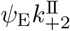 via the combined action of concentration partitioning (*ζ*_X_) and kinetic partitioning (*ζ*_*β*_), reflecting the competition between concentration enrichment of enzyme-bound complexes and kinetic re-weighting of the catalytic step.

##### Thermodynamic contribution

The thermodynamic contribution 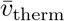 can be expressed as

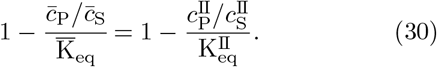

Because this term is directly related to the reaction free energy Δ*µ* = *µ*_P_ −*µ*_S_ (see Sec. II C of the Supplementary Information), Eq. (30) shows that phase separation leaves the thermodynamic driving force unchanged relative to the reference homogeneous system.

##### Binding contribution

The enzyme-binding contribution 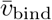 depends on both the redistribution of clients and the kinetic re-weighting of the underlying reaction steps. This dependence is encoded in the systemaveraged Michaelis constants 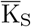 and 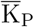, which are non-linear combinations of the system-averaged rate constants 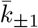 and 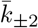 (Eq. (27)). Consequently, they do not admit a simple volume-weighted representation in terms of phase-specific Michaelis constants; instead,

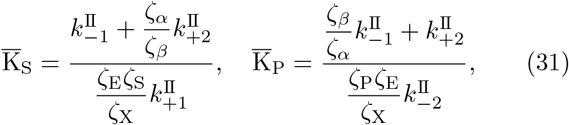

for which the combinations 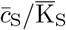 and 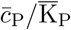 appearing in the saturation term become

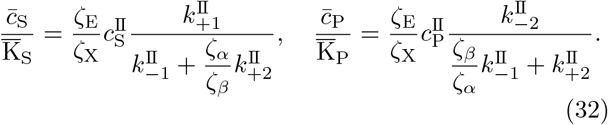

These expressions show that, unlike the thermodynamic contribution, the enzyme-binding contribution 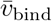 is jointly regulated by concentration partitioning and kinetic partitioning. As a result, phase separation modifies enzyme saturation through both redistribution of free and bound enzyme species and through kinetic re-weighting of the reaction steps.

##### Physical interpretation

Taken together, the kinetic and enzyme-binding contributions emerge as the only sources of condensate-mediated regulation under reaction-limited conditions, entirely controlled by concentration partitioning and kinetic partitioning. While the thermodynamic driving force remains unchanged, these two mechanisms jointly renormalize the kinetic capacity and enzyme saturation of the effective homogeneous reaction scheme (Eqs. (29) and (32)) through the redistribution of client species between phases and the re-weighting of microscopic reaction kinetics.

The homogeneous reference system is recovered in the limit V^I^*/*V → 0, for which all concentration and kinetic partitioning degrees approach unity (*ζ*_*i*_ → 1, *ζ*_*η*_ → 1). In this limit, the renormalized kinetic and binding contributions reduce to their homogeneous values, and Eq. (28) recovers the standard reversible Michaelis–Menten rate law for homogeneous systems derived in the Supplementary Information (cf. Eq. (S11)).

Eq. (28) provides the system-averaged initial rate 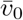 in the reaction-limited regime (Fig. 4). In Fig. 4a, it corresponds to the upper curve (*ℓ*_min_ ≫ *R, L*), which exhibits the largest enhancement relative to the homogeneous reference system. This large response arises because transport rapidly redistributes clients across phases, continuously replenishing reactants within the condensate and allowing condensate-localized kinetic enhancement to be fully expressed at the system level. Evaluating Eq. (28) at the initial substrate concentration highlighted in Fig. 4a yields the reaction-limited asymptote identified by the square marker.

#### d. Relative initial rate

The relative initial rate ℛ, introduced in Sec. III A (Eq. (15)) as a numerical measure of condensate-mediated regulation, admits a useful analytical factorization under reaction-limited conditions.

Using the factorized expression for the reaction-limited initial rate (Eq. (28)) and its homogeneous limit obtained by setting *ζ*_*i*_ → 1 and *ζ*_*η*_ → 1 (Eq. (S11), Supplementary material), the relative initial rate factorizes into kinetic and enzyme-binding contributions (the thermodynamic contribution cancels identically because it is the same in phase-separated and homogeneous systems, Eq. (30)),

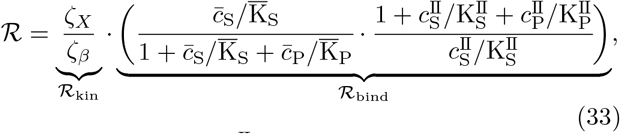

where 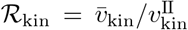 and 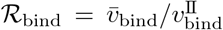. Unless otherwise stated, concentrations appearing in the initial-rate expressions are evaluated at the quasi–steadystate time used to define the corresponding initial rate. The factorization in Eq. (33) makes explicit how concentration and kinetic partitioning jointly regulate ℛ through the partitioning degrees of free and bound enzyme species, *ζ*_E_ and *ζ*_X_, and the kinetic partitioning degrees associated with the binding and catalytic steps, *ζ*_*α*_ and *ζ*_*β*_. Condensate size V^I^ enters implicitly through these quantities. Eq. (33) thus provides the analytical framework used below to study the conditions for optimal enhancement and suppression of catalytic activity.

### Conditions for optimal regulation of ℛ

To characterize the emergence of optimal condensate sizes regulating catalytic activity, we consider extrema of the relative initial rate with respect to the condensate volume fraction *ϕ* ≡ V^I^*/*V for fixed client partitioning and kinetic partitioning. These extrema are defined by 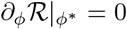. For regimes in which ℛ develops at most a single extremum, the existence of such an extremum is completely determined by the signs of the small-condensate and large-condensate asymptotic slopes.

We therefore study the sign of ∂_*ϕ*_ ℛ in these limits, which provides the necessary and sufficient conditions for the existence of the extremum. Since ℛ *>* 0, the derivatives ∂_*ϕ*_ ℛ and ∂_*ϕ*_ ln ℛ always have the same sign and vanish at the same value of *ϕ*. Extrema of can ℛ therefore be identified equivalently from the logarithmic slope ∂_*ϕ*_ ln ℛ, which admits a simpler analytical form (expressions are given in Appendix E).

#### Enzyme-substrate co-localization regime

When enzyme and substrate co-localize within the condensate (*P*_E_ *>* 1, *P*_S_ *>* 1; cf. Fig. 5), ℛ develops at most a single maximum over the explored parameter range. The existence of an optimal condensate size therefore requires a positive small-condensate asymptotic slope and a negative large-condensate asymptotic slope, equivalently

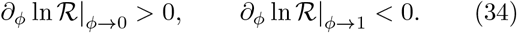

These conditions correspond directly to the small-condensate and large-condensate limits used to interpret the emergence of optimal enhancement in Sec. III. For the parameter regime considered here, the large-condensate slope ∂_*ϕ*_ ln ℛ |_*ϕ*→1_ remains negative throughout the explored range of catalytic-step kinetic partition coefficients (see Appendix E). Consequently, the emergence of an optimum is controlled entirely by the sign of the small-condensate slope ∂_*ϕ*_ ln ℛ |_*ϕ*→0_, yielding the critical catalytic-step kinetic partition coefficient

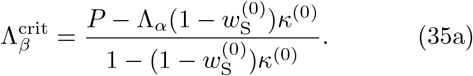

Here, 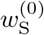 denotes the substrate saturation weight of the homogeneous reference system, whereas *κ*^(0)^ quantifies the probability that a bound complex proceeds to catalysis rather than dissociation (Appendix E). For 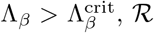, ℛ develops a maximum at finite condensate size; for 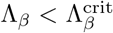, it remains monotonic. This analytical criterion explains the transition observed in Fig. 5a, where 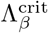 separates monotonic suppression from the emergence of optimal enhancement.

In the regime when complex dissociation dominates over catalytic turnover 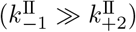, the correction term 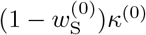 is small, yielding

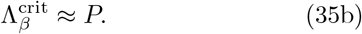

In the limit 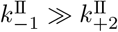, the simple relation in Eq. (35b) shows that the onset of optimality is determined by a direct competition between catalytic enhancement and client enrichment, consistent with the physical interpretation developed in Sec. III and illustrated in Fig. 5.

#### Enzyme-substrate spatial segregation

When the enzyme is excluded from the substrate-enriched condensate (*P*_E_ *<* 1, *P*_S_ *>* 1; cf. Fig. 6), the relative initial rate can exhibit qualitatively distinct behaviours depending on the catalytic-step kinetic partition coefficient: monotonic regulation (no extrema), optimal suppression (a single minimum), or optimal enhancement (a minimum and a maximum). We do not pursue a complete analytical characterization of this regime here because, unlike the enzyme-substrate co-localization regime, spatial segregation can produce multiple extrema of ℛ. Consequently, the signs of the small-condensate and large-condensate asymptotic slopes are no longer sufficient to determine the existence of optimal suppression or optimal enhancement. Nevertheless, the asymptotic slopes remain useful as they provide intuition for interpreting the richer regulatory landscape observed in Fig. 6.

## IV. CONCLUSION

Membrane-less condensates formed through liquidliquid phase separation are increasingly recognized as important regulators of biomolecular catalysis in synthetic and living cells [18–20]. Insights from studies in cells, complemented by *in vitro* studies, have shown that condensates can regulate enzymatic activity by enhancing reaction rates, redirecting metabolites between competing pathways according to cellular demand, and switching enzyme activity on and off through reversible compartmentalization [24, 25, 75–77]. Biochemical mechanisms underlying such regulatory effects are based on changes in the local physicochemical environment within condensates, including reactant enrichment, reactant sequestration, altered molecular transport, and modifications of reaction kinetics [22–26, 78]. Yet a general quantitative framework capable of predicting how phase separation controls catalysis has remained largely lacking [27–30].

### A quantitative framework for condensate-mediated catalysis

In this work, we provide such a framework and show that condensate-mediated catalysis cannot, in general, be understood from molecular enrichment alone. Instead, catalytic regulation emerges from the coupled effects of molecular partitioning, diffusive transport, and phase-dependent reaction kinetics. Building on recent theoretical studies which started to unravel the physicochemical principles underlying condensate-mediated regulation of chemical reactions [35, 79], we establish a quantitative and thermodynamically consistent theory of biomolecular catalysis in phase-separated systems based on reversible Michaelis–Menten kinetics. The resulting framework links condensate properties directly to catalytic readouts and is formulated entirely in terms of experimentally relevant parameters and measurable quantities, including condensate size, molecular partition coefficients, system-averaged concentrations, diffusion coefficients, and reaction rate constants.

### Transport as a regulator of catalytic response

A central finding of our work is that condensate-mediated regulation cannot, in general, be understood from molecular enrichment alone. Instead, condensate-mediated catalysis emerges from the coupled action of concentration partitioning, reaction kinetic partitioning, and diffusive transport. Phase separation not only selectively redistributes enzymes and substrates between coexisting phases, but also modifies the effective kinetics of individual reaction steps, while transport couples the resulting reaction dynamics by mediating the exchange of reacting species across phase boundaries. Together, these mechanisms determine whether condensates enhance or suppress catalytic activity and how strongly condensate-localized kinetics is expressed at the system level.

Diffusive transport emerges as a key regulator of this process. Depending on condensate size, molecular partitioning of reactants, and reaction kinetic partitioning, slow interphase exchange confines kinetic advantages to individual phases, whereas rapid exchange allows condensate-localized kinetics to be fully expressed at the system level. Remarkably, under rapid exchange conditions, the spatially heterogeneous catalytic system admits a coarse-grained description in terms of system-averaged concentrations and reaction kinetics. This reduction transforms the complexity of phase-separated reaction-diffusion dynamics into a predictive kinetic theory, yielding explicit analytical conditions for the rich landscape of condensate-mediated regulation.

Accordingly, we find the strongest effects of condensate-mediated regulation occurring in the regime of rapid exchange, where *µ*m-sized condensates can either enhance catalytic activity or suppress it relative to systems without condensates. In particular, for the experimentally motivated parameter set considered here, enzyme-substrate co-localization conditions give rise to optimal enhancement of catalytic activity by up to 100fold within a relatively narrow range of condensate sizes (*R* ≈ [0.6, 3] *µ*m). By contrast, enzyme-substrate spatial segregation conditions yield a richer phenomenology, including monotonic enhancement and suppression, optimal enhancement by up to two orders of magnitude, and optimal suppression approaching one order of magnitude, with the emergence of optimal condensate sizes over a broader range (*R* ≈ [1, 6] *µ*m for optimal enhancement; *R* ≈ [4, 10] *µ*m for optimal suppression). Notably, the predicted condensate radii mediating optimal regulation fall within the broad micron-scale range reported for many biomolecular condensates. Importantly, these results demonstrate that spatial segregation does not necessarily impair catalytic performance. Rather, enzyme depletion and kinetic enhancement cooperate to generate a rich spectrum of regulatory behaviours, with sufficiently strong kinetic enhancement more than compensating for the reduced enzyme abundance within the condensate.

### Biological implications and experimental validation

These findings provide a quantitative framework for understanding the diverse regulatory effects observed in biomolecular condensates [24, 25, 78], ranging from G-body-mediated enhancement of glycolysis in living cells [78] to engineered condensates displaying up to 36-fold increase in SUMOylation enzyme cascade activity [29]. Moreover, they reveal that neither local reactant enrichment nor accelerated kinetics is sufficient, by itself, to predict catalytic response. From a broader perspective, the presence of both enhancement and suppression within the same framework supports the view that condensates are not universal amplifiers of catalysis nor passive concentrating devices, but rather tunable reaction environments which can act as catalytic switches. By tuning condensate size, partitioning, or transport properties, phase separation can move a reaction network between regimes of enhanced and suppressed activity, thus providing a powerful mechanism for regulation [78].

Beyond individual enzymatic reactions, the ability of condensates to switch catalytic activity between enhanced and suppressed states suggests a general mechanism through which cells may dynamically redirect metabolic fluxes and reorganize biochemical pathways in response to changing physiological demands. To test the predicted existence of optimal condensate sizes and transitions between enhancement and suppression regimes, we propose reconstituted phase-separated enzymatic systems in which condensate size and molecular partitioning can be independently controlled. Particularly promising are recently developed condensates based on intrinsically disordered proteins, including systems involving adenylate kinase and phase-separating DEAD-box proteins [80, 81], where condensate formation was shown to significantly increase (60to 100-fold) enzyme concentrations within the condensate as compared to the starting concentration, while simultaneously altering transport properties (25to 200-fold increase in viscosity in the condensate) and catalytic activity (5-fold increase in initial velocity when compared to a reaction in homogeneous solution). The key observables predicted by our theory, including partition coefficients, concentrations, diffusivities, initial rates, and half-times, are directly accessible through fluorescence microscopy, activity assays, and emerging label-free spectroscopic techniques.

More broadly, our results show that biomolecular condensates are not passive concentrating devices whose effects can be inferred from reactant enrichment alone. Rather, they act as tunable reaction environments in which partitioning, transport, and reaction kinetics collectively control catalytic activity. This perspective provides a quantitative foundation for understanding, predicting, and engineering condensate-mediated catalysis.

## ACKNOWLEDGMENTS

We thank H. Vuijk and A. Thatte for valuable feedback on the manuscript, and D. Tang, A. Gosh and M. Gao for fruitful discussions on enzymatic reaction in the PEN-DNA system. The authors used Microsoft 365 Copilot (GPT-5-based model) to assist with language editing and text revision. C. Weber acknowledges the European Research Council (ERC) under the European Union’s Horizon 2020 research and innovation program (Fuelled Life, Grant No. 949021).

## Appendix A Partitioning degree

### a. Concentration partitioning degrees ζ_i_(P_i_, V^I^)

The distinct physicochemical environments imposed by the scaffold redistribute client species between coexisting phases. For a species *i*, phase equilibrium sets its relative enrichment or depletion between the two phases through the partition coefficient (Eq. (2))

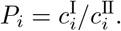

The values 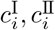 are not independent, but are constrained by conservation of total material (Eq. (21a)),

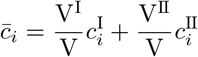

where 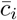 denotes the system-averaged concentration. Solving these relations yields Eq. (21b),

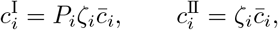

with

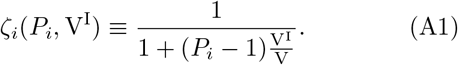

The quantity *ζ*_*i*_ is the *partitioning degree* of client species *i*. It measures how the system-averaged concentration 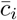 is distributed between the two phases for a given partition coefficient *P*_*i*_ and condensate size V^I^. Equivalently, 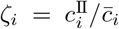 quantifies the fraction of the system-averaged concentration carried by phase II. The partitioning degree therefore maps the phase-specific concentrations with their system-level representation.

#### a. Limiting cases

The partitioning degree *ζ*_*i*_ continuously interpolates between two limiting regimes. To illustrate their physical meaning, we consider the case *P*_*i*_ *>* 1, corresponding to a client species enriched in the condensate relative to the dilute phase.

#### Small-condensate limit

When *ϕ* ≡ V^I^*/*V → 0,

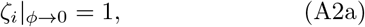

such that

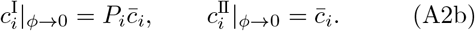

In this limit, the condensate occupies a vanishing fraction of the system volume. Therefore, even though the concentration of client species *i* within the condensate satisfies 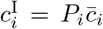, its contribution to the system-averaged concentration becomes negligible. The system-averaged concentration is therefore determined entirely by the dilute phase, 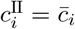, independent of the value of *P*_*i*_. Consequently, the system becomes effectively indistinguishable from a homogeneous phase-II system.

#### Large-condensate limit

Conversely, when *ϕ* → 1,

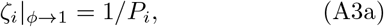

such that

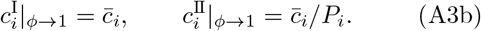

In this limit, the condensate occupies nearly the entire system volume. Consequently, the system-averaged concentration becomes entirely determined by the condensate phase, 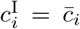. The partition coefficient *P*_*i*_ is still satisfied through 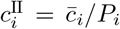, but the dilute phase occupies a negligible fraction of the system volume and therefore has a negligible influence on the system average. As a result, the system becomes effectively indistinguishable from a homogeneous phase-I system.

#### Intermediate condensate sizes

For 0 *< ϕ <* 1, the partitioning degree *ζ*_*i*_ determines how the total amount of client species *i* is distributed between the two phases. As the condensate volume fraction *ϕ* increases, partitioning becomes progressively less effective at generating enrichment relative to the system average. Specifically, the enrichment of species *i* in the condensate relative to the system average,

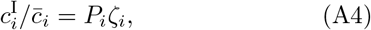

decreases continuously from *P*_*i*_ (at *ϕ* → 0; Eq. (A2)) to unity (at *ϕ* → 1; Eq. (A3)). Thus, species enriched within the condensate (*P*_*i*_ *>* 1) become progressively less enriched as the condensate occupies a larger fraction of the system volume, whereas species depleted from the condensate (*P*_*i*_ *<* 1) become progressively less depleted.

### b. Kinetic partitioning degrees ζ_η_ (Λ_η_, V^I^)

In addition to redistributing client concentrations, the distinct physicochemical environments imposed by the scaffold can modify the microscopic kinetics of individual reaction steps. This effect is captured by the phasedependent kinetic prefactors 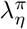, whose values may differ between the coexisting phases. Unlike concentrations 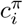, however, kinetic prefactors are neither conserved quantities nor constrained by phase equilibrium. Consequently, no conservation relation analogous to Eq. (21a) exists for the kinetic prefactors themselves. Nevertheless, the *kinetic partitioning degree ζ*_*η*_ emerges from the reactionlimited coarse-graining of the phase-resolved reaction kinetics (see Appendix D),

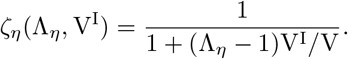

Although mathematically equivalent, *ζ*_*i*_ and *ζ*_*η*_ have fundamentally different physical origins. Whereas *ζ*_*i*_ follows from phase equilibrium and conservation of total material, and quantifies how client species contribute to the system-averaged concentrations through their redistribution between phases, the kinetic partitioning degree *ζ*_*η*_ originates from the kinetic coarse-graining under reaction-limited conditions, and quantifies how phase-dependent microscopic kinetics of reaction step *η* contribute to the effective system-averaged rate constants.

#### a. Limiting cases

Despite their different physical origins, *ζ*_*i*_ and *ζ*_*η*_ share the same mathematical form and therefore obey the same limiting behaviour with respect to condensate size. The physical meaning of these limits, however, is fundamentally different.

##### Small-condensate limit

When *ϕ* ≡ V^I^*/*V → 0,

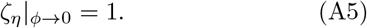

To illustrate the physical meaning of this limit, consider the system-averaged rate constant associated with the forward contribution of reaction step *β* (Eq. (27b)),

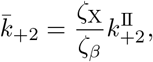

which follows from the reaction-limited coarse-graining of the phase-resolved kinetics (Appendix D). As *ϕ* → 0, both *ζ*_X_ and *ζ*_*β*_ approach unity, yielding

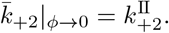

The effective system-level kinetics therefore reduces to that of the homogeneous phase-II system.

##### Large-condensate limit

Conversely, when *ϕ* → 1,

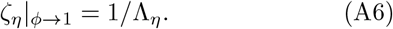

Consider again the representative system-averaged rate constant associated with the forward contribution of reaction step *β*,

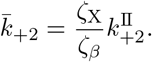

Using Eqs. (A3a) and (A6), one obtains

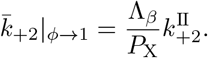

Thus, both concentration partitioning and kinetic partitioning remain explicitly encoded in the system-averaged rate constant even when the condensate occupies nearly the entire system volume. However, whereas the enrichment generated by concentration partitioning vanishes as *ϕ* → 1 (Eq. (A4)), kinetic partitioning continues to directly modulate the effective reaction kinetics through its explicit contribution to the system-averaged rate constants. The kinetic effects of the condensate persist even in the large-condensate limit.

## Appendix B Half-time in phase-separated systems

While the initial rate characterizes the early-time catalytic response near the onset of quasi–steady state, it restricts the analysis to a narrow temporal window and discards information contained in the subsequent evolution of the system [41]. A complementary observable is the *reaction half-time*, which provides a global measure of the progress of catalysis toward equilibrium and is particularly useful when initial-rate measurements are difficult or catalysis approaches saturation [82].

For reversible enzymatic reactions, two natural definitions can be considered: a product half-formation time, 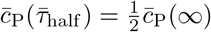, and a substrate half-depletion time, 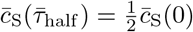. These definitions coincide in the irreversible limit, where the reaction proceeds essentially to completion, 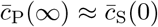. They generally differ for reversible reactions because complete substrate-to-product conversion is not achieved at equilibrium.

Throughout this work, we adopt the substrate half-depletion time, consistent with the classical integrated-rate analysis of Alberty and Koerber [83],

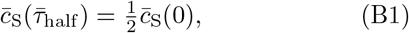

i.e., the time required for the system-averaged substrate concentration to decrease to half of its initial value. This definition is used throughout all results presented below.

### a. Transport-controlled reduction of the half-time

We first examine how the half-time varies with the initial substrate concentration across the kinetic regimes, while keeping all condensate properties fixed (Table I). For direct comparison with the enzyme-normalized initial rate 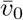 (cf. Fig. 4a), we plot the inverse half-time 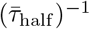, so that both observables have units of inverse time and quantify the rate of catalytic progression. The dependence of the inverse half-time on the kinetic regime closely parallels that observed for 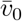. For the parameter set considered here, condensates increase the inverse half-time relative to the homogeneous reference system across all initial substrate concentrations explored, thereby reducing the half-time and accelerating the approach to equilibrium (Fig. 7). The magnitude of this enhancement increases continuously from diffusion-limited to reaction-limited behaviour, indicating that transport controls not only the initial rate but also the half-time.

**Figure 7.**
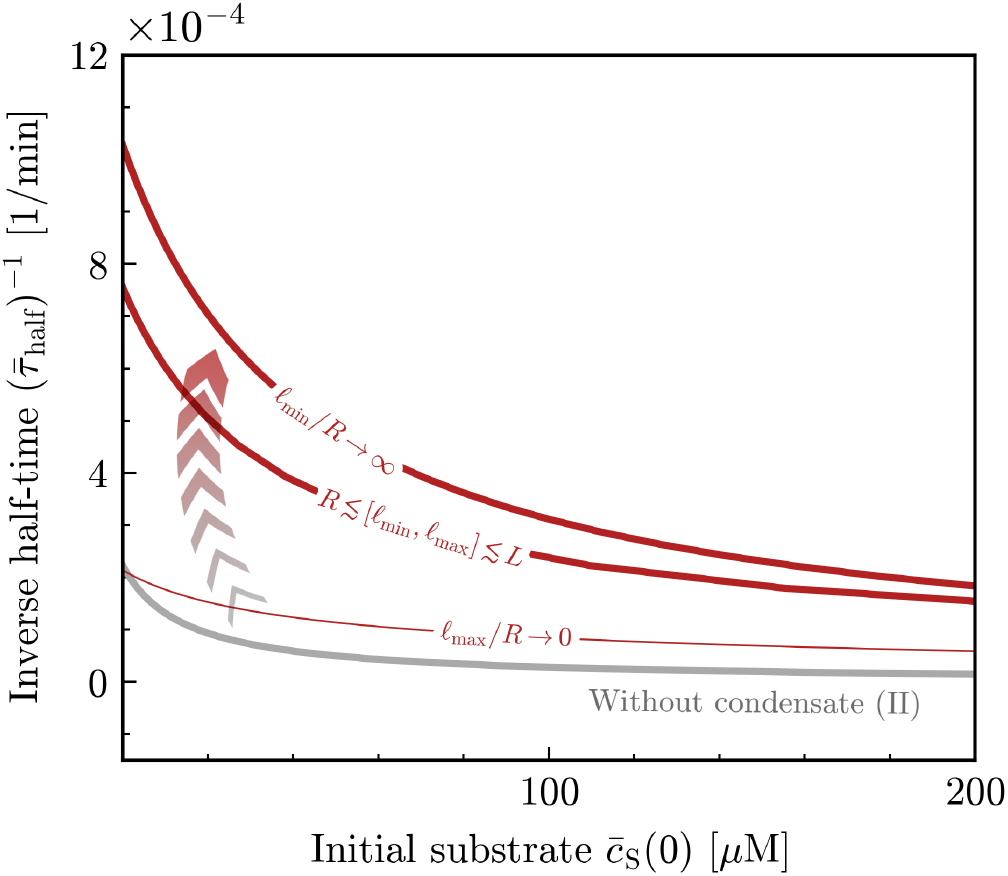
Transport-controlled reduction of the substrate half-time mediated by condensates. Inverse halftime 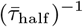 as a function of the initial substrate concentration 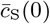, comparing systems with a phase-separated conden-sate (red) to the homogeneous reference system without condensates (gray). Red curves correspond to diffusion-limited, reaction–diffusion, and reaction-limited behaviour (cf. Fig. 4). For the diffusion-limited asymptotic limit (*ℓ*_max_*/R* → 0), the reported half-time is constructed from a volume-weighted average of the phase-specific half-times and should be interpreted as a descriptive coarse-grained proxy rather than a unique collective timescale. Parameter values in Table I.

**Figure 8.**
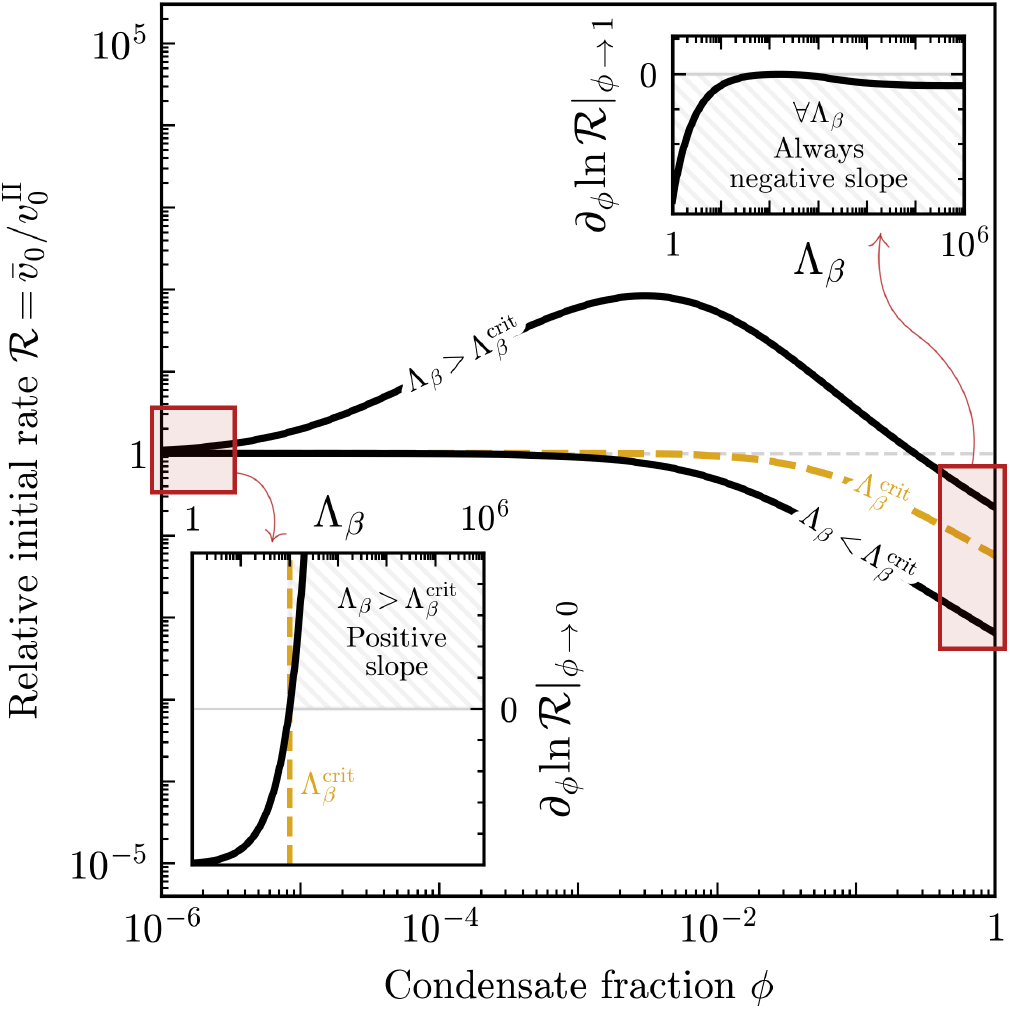
Asymptotic slope conditions for condensatemediated optimal enhancement in the enzyme-substrate co-localization regime. Representative condensate-size dependence of the relative initial rate ℛ for kinetic partition coefficients above, below, and at the critical value 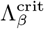. The emergence of a maximum in ℛ is governed by the asymptotic slopes at small and large condensate fractions. **Insets**: the small-condensate asymptotic slope ∂_*ϕ*_ ln ℛ|_0_ (Eq. (E6)) changes sign at the critical value 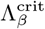, whereas the large-condensate asymptotic slope ∂_*ϕ*_ ln ℛ|_1_ (Eq. (E7)) remains negative throughout the explored parameter range. Consequently, optimal enhancement emerges when 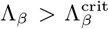, for which ∂_*ϕ*_ ln ℛ|_0_ *>* 0 and ∂_*ϕ*_ ln _1_ *<* 0. Red boxes indicate the asymptotic regions analysed by the corresponding insets. For parameter values, see Table I.

Closed-form expressions for the half-time are obtained in the asymptotic limits *ℓ*_max_*/R* → 0 and *ℓ*_min_*/R* → ∞ of the diffusionand reaction-limited regimes. These expressions follow from the homogeneous Michaelis– Menten solution derived in the Supplementary Information and are evaluated using the corresponding phasespecific or coarse-grained kinetic parameters.

### b. Diffusion-limited kinetics

In the asymptotic limit *ℓ*_max_*/R* → 0, the two phases evolve as kinetically isolated compartments governed by phase-specific kinetics. Because phases evolve independently, there is no meaningful system-level half-time associated with the phase-separated system. The only well-defined half-times are the phase-specific quantities 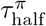, defined by

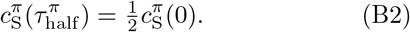

The phase-specific half-times are therefore obtained by integrating the corresponding phase-specific Michaelis– Menten rate law (Eq. (19)) and applying Eq. (B2). The resulting expression follows directly from the homogeneous half-time solution derived in Sec. II C of the Supplementary Information, evaluated with the corresponding phase-specific kinetic parameters,

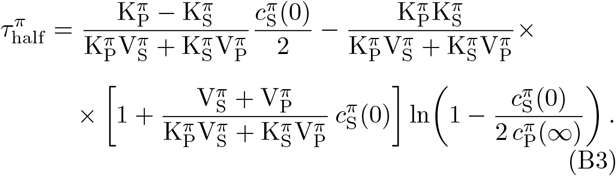

Here, 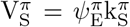 and 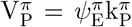 denote the maximal forward and reverse reaction velocities, respectively. Eq. (B3) provides the phase-specific half-times 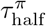 in the diffusion-limited asymptotic limit (*ℓ*_max_*/R* → 0; Fig. 7: for graphical comparison with the other regimes, the diffusion-limited curve 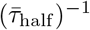 is constructed from a volume-weighted average of the phase-specific half-times. This quantity should be regarded as a descriptive coarsegrained proxy rather than a unique system-level relaxation timescale.

### c. Reaction-limited kinetics

In the asymptotic limit *ℓ*_min_*/R* → ∞, rapid interphase diffusion maintains phase equilibrium throughout the system. Consequently, client concentrations evolve on a common global timescale, and the system-level half-time 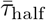 becomes a meaningful observable of the coarse-grained reaction scheme.

Integrating the coarse-grained Michaelis–Menten rate law (Eq. (28)) and applying the half-depletion condition (Eq. (B1)) yields the homogeneous half-time expression derived in Sec. II C of the Supplementary Information, evaluated using the coarse-grained kinetic parameters,

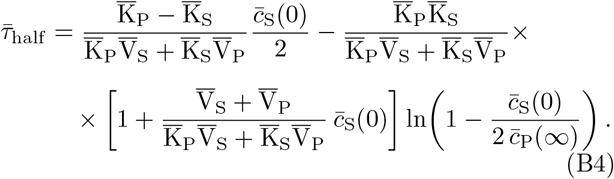

Here, 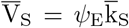 and 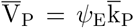 denote the maximal forward and reverse reaction velocities, respectively. Consistent with the system-level coarse-graining available in the reaction-limited asymptotic limit, the halftime is entirely determined by the renormalized kinetic parameters of the coarse-grained reaction scheme and is therefore associated with a single global kinetic timescale. Eq. (B4) provides the system-level half-time 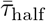 in the reaction-limited asymptotic limit (*ℓ*_min_*/R* → ∞; Fig. 7).

## Appendix C Reaction–diffusion length scales

Reaction–diffusion length scales quantify the spatial extent over which concentration perturbations spread by diffusion before being significantly altered by reactions. They provide a convenient metric to characterize the interplay between diffusive transport and reaction kinetics.

### Formal definition

Reaction–diffusion length scales are determined from the eigenmodes of the linearized reaction–diffusion dynamics. For the reversible Michaelis–Menten kinetics, characterized by non-linear reaction terms (both first-order and second-order processes are present), this would require linearization of a four-species coupled reaction–diffusion system (Eq. (1)) followed by analysis of the resulting eigenmodes.

### Practical construction

Rather than explicitly analysing these eigenmodes, we introduce reaction– diffusion length-scale estimates that provide order-of-magnitude measures of the competition between diffusion and the individual reaction steps. To construct these estimates, the non-linear bimolecular terms (enzyme– substrate association in step *α* and product rebinding in step *β*) are locally linearized around a reference concentration 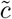 (e.g., an initial, steady-state, or representative concentration of the partner species). Because these quantities are intended only as order-of-magnitude estimates, the precise choice of the reference concentration 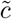 affects the resulting values only quantitatively and is not expected to alter the qualitative regime classification. This yields effective first-order rate constants

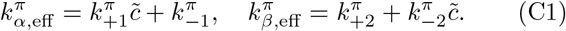

The associated reaction–diffusion length scale estimate 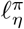 for reaction step *η* in phase *π* is

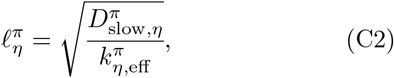

where 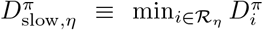 is the slowest diffusivity among the species involved in reaction step *η* (e.g., ℛ_*α*_ = E, S, X and ℛ_*β*_ = X, E, P ). The slowest diffusivity provides a simple and conservative estimate of the distance over which concentration perturbations can propagate, while the effective rate constant sets the characteristic rate at which they are relaxed by reactions.

### Kinetic-regime indicators

The smallest and largest of these length scales,

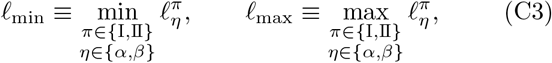

provide lower and upper bounds on the reaction–diffusion length scales associated with the individual steps of the Michaelis–Menten reaction kinetics and therefore serve as useful indicators for the transition between reaction-limited and diffusion-limited behaviour (cf. Fig. 3).

Because 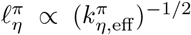, the smallest length scale *ℓ*_min_ is set by the fastest effective reaction relative to diffusion (i.e., the largest 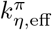, given the limiting diffusivity 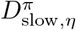 ). Conversely, the largest length scale *ℓ*_max_ is set by the slowest effective reaction relative to diffusion (i.e., the smallest 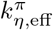, given the limiting diffusivity 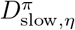). These two limits respectively characterize the most and least efficient suppression of concentration gradients.

## Appendix D Reaction-limited regime: from phase-specific apparent rate constants to system-averaged rate constants

Under reaction-limited conditions, the kinetic coarsegraining introduced in the main text proceeds in two steps. First, the phase-resolved reaction fluxes 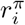 are rewritten in terms of system-averaged concentrations, which introduces phase-specific apparent rate constants 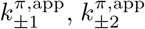. Second, these apparent rate constants are combined through volume-weighted averaging to yield the system-averaged rate constants 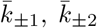 appearing in the coarse-grained reaction scheme (Eq. (26)).

### a. Phase-specific apparent rate constants

#### Forward step α (representative)

The first step of the kinetic coarse-graining consists of expressing the phase-resolved reaction fluxes in terms of the system-averaged concentrations 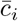. To illustrate this construction, we consider the forward contribution of reaction step *α*. The corresponding phase-specific effective rate constants 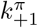 are given by Eq. (6c). Substituting the phase equilibrium conditions (Eq. (21b)) in the forward flux 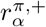 yields

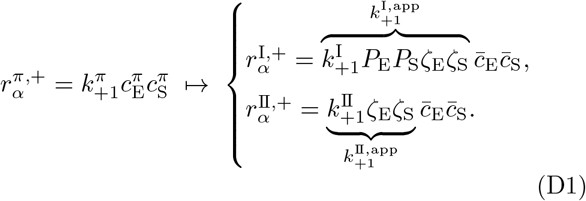

This motivates the introduction of the phase-specific apparent rate constants 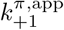, defined through

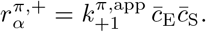

The term *apparent* emphasizes that these quantities are not intrinsic kinetic parameters of the individual phases. Rather, they emerge from rewriting local reaction fluxes in terms of the global dynamical variables 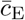 and 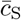 under the constraint of phase equilibrium.

Using the relations between the effective rate constants in the two phases, which determine the condensate value 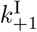 from the homogeneous value 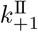 (Eq. (6c)), one finds

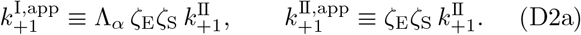

The phase-specific apparent rate constants associated with the forward contribution of reaction step *α* are therefore determined by two distinct effects. The factor *ζ*_E_*ζ*_S_ reflects the redistribution of the species participating in the reaction step and therefore contributes to the apparent kinetics in both phases. In contrast, the kinetic partition coefficient Λ_*α*_ captures the condensate-induced modification of the microscopic reaction kinetics of the step, and contributes only to the condensate apparent rate constant 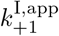. The apparent rate constants thus constitute the phase-specific kinetic ingredients entering the reaction-limited coarse-graining.

#### Other reaction steps

The same construction can be applied to the remaining reaction steps, yielding

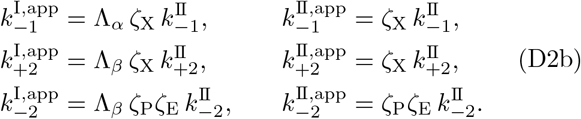

### b. System-averaged rate constants

#### Forward step α (representative)

The apparent rate constants introduced in Eq. (D2) are the basis of the kinetic coarse-graining. Analogously to the construction of the system-averaged concentrations and reaction fluxes, the second step consists of combining the phase-specific apparent rate constants through volume-weighted averaging to obtain the system-averaged rate constants appearing in the effective homogeneous reaction scheme.

For the forward contribution of reaction step *α*, the system-averaged rate constant 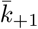 is obtained by volume-weighted averaging of the phase-specific apparent rate constants,

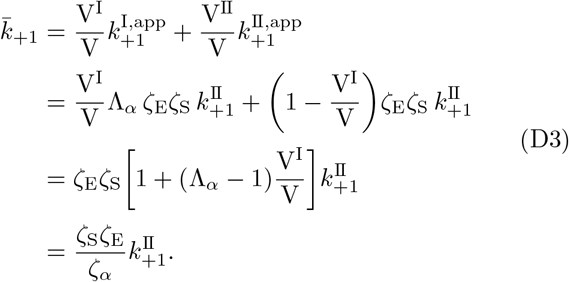

The final expression naturally introduces the kinetic partitioning degree *ζ*_*α*_(Λ_*α*_, V^I^) discussed in Appendix A. Importantly, whereas the concentration partitioning degrees *ζ*_*i*_ act directly on the phase concentrations and therefore govern the enrichment of client species relative to the system average, the kinetic partitioning degree *ζ*_*α*_ acts on the coarse-grained rate constants themselves and therefore governs how phase-specific microscopic kinetics contribute to the system-level description.

#### Other reaction steps

Applying the same construction to the remaining reaction steps yields the system-averaged rate constants presented in the main text (Eq. (27)).

## Appendix E Conditions for optimal regulation of the initial rate under reaction-limited kinetics

This appendix derives the asymptotic slopes of the relative initial rate ℛ in the limits *ϕ* ≡ V^I^*/*V → 0 and *ϕ* → 1 discussed in Secs. III. These slopes provide the analytical conditions used to characterize the emergence of optimal enhancement and suppression of catalytic activity.

### a. Small-condensate expansion of ℛ

In the smallcondensate limit, *ϕ* ≡ V^I^*/*V → 0, all concentration and kinetic partitioning degrees, *ζ*_*i*_ and *ζ*_*η*_, approach unity. As discussed in Appendix A, the phase-separated system in this limit becomes indistinguishable from a homogeneous phase-II system, yielding

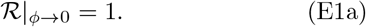

Expanding the relative initial rate around *ϕ* = 0 gives

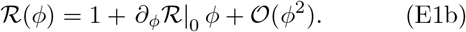

The leading-order effect of condensate formation is therefore entirely determined by the sign of ∂_*ϕ*_ ℛ|_0_, i.e., the small-condensate slope. Positive values indicate that ℛ increases as a condensate forms, whereas negative values indicate that it decreases. The small-condensate slope therefore provides the first indication of whether condensate formation enhances or suppresses ℛ relative to the homogeneous reference system. However, the existence of extrema in ℛ (and therefore of optimal condensate-mediated regulation) depends on its behaviour across the full range of condensate sizes. Although the analysis of the asymptotic slopes presented below does not capture all possible behaviours of ℛ, it provides sufficient conditions for the existence of an extremum, defined by ∂_*ϕ*_ ℛ = 0, whenever ℛ exhibits at most a single extremum as a function of *ϕ*.

### b. Asymptotic slopes

Since ℛ *>* 0, from ∂_*ϕ*_ ℛ = ℛ *·* ∂_*ϕ*_ ln ℛ we see that the derivatives ∂_*ϕ*_ and ∂_*ϕ*_ ln have the same sign. We therefore analyse the logarithmic slope ∂_*ϕ*_ ln ℛ instead, as this considerably simplifies the analysis. Because the relative initial rate factorizes into kinetic and enzyme-binding contributions (Eq. (33)),

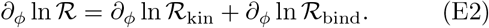

Accordingly, the response of both the concentration and kinetic partitioning degrees, *ζ*_*i*_ and *ζ*_*η*_, to changes in condensate size can be characterized through their logarithmic derivatives. For any partitioning degree *ζ*(*X, ϕ*), one obtains

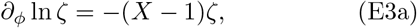

which in the asymptotic limits *ϕ* → 0 and *ϕ* → 1 simplifies to

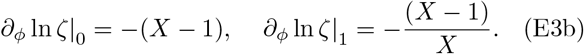

Eqs. (E2) and (E3) provide the key ingredients required for the analytical evaluation of the asymptotic slopes and, in regimes where ℛ exhibits at most a single extremum, for deriving conditions for its existence.

#### Small-condensate limit

As discussed above, the sign of the small-condensate slope ∂_*ϕ*_ ℛ|_0_ determines whether the formation of a condensate increases or decreases ℛ. The corresponding asymptotic logarithmic slope reads

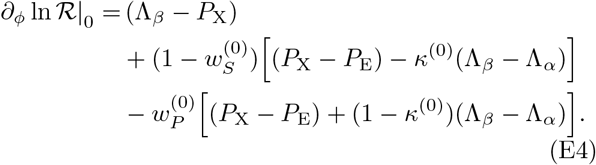

The first term, (Λ_*β*_ *−P*_X_), captures the competition between kinetic partitioning of the catalytic step and equilibrium partitioning of the complex. The remaining terms arise from the enzyme-binding contribution, where different combinations of partition coefficients are weighted by

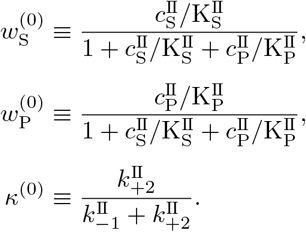

Here, 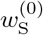 and 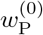 represent the substrate and product saturation weights of the homogeneous reference system, whereas *κ*^(0)^ measures the catalytic *commitment* of the complex, i.e., the probability that a bound complex proceeds to catalysis rather than dissociation.

#### Large-condensate limit

In the large-condensate limit (*ϕ* → 1), the corresponding asymptotic logarithmic slope becomes

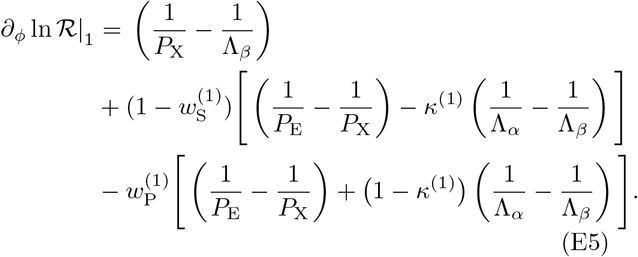

As for the small-condensate limit, the first term reflects the competition between equilibrium partitioning of the complex and kinetic partitioning of the catalytic step, while the remaining terms arise from the enzyme-binding contribution and are modulated by

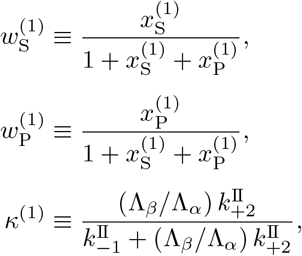

with

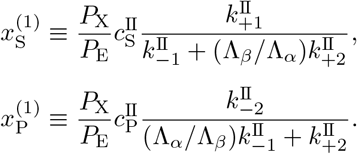

### c. Enzyme-substrate co-localization regime

To connect the general asymptotic conditions derived above with the results presented in Sec. III, we now specialize the logarithmic slopes in Eqs. (E4) and (E5) to the enzyme–substrate co-localization regime (*P*_*E*_ *>* 1, *P*_*S*_ *>* 1). Consistent with the parameter choices used throughout this work, we assume equal client partition coefficients, *P*_E_ = *P*_S_ = *P*_P_ = *P*_X_ ≡ *P*, together with Λ_*β*_ *>* Λ_*α*_ *>* 1 and the initial conditions 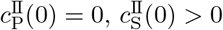.

Under these assumptions, product-dependent contributions vanish 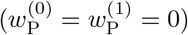 and the dependence on differential client partitioning cancels (*P*_E_ − *P*_X_ = 0). The asymptotic slopes therefore depend only on the competition between concentration partitioning (*P* ) and kinetic partitioning (Λ_*α*_ and Λ_*β*_), modulated by the substrate-saturation and kinetic weights 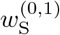 and 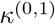.

The asymptotic slope at *ϕ* → 0 (Eq. (E4)) reduces to

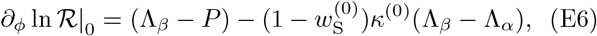

where

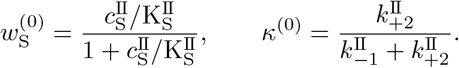

Similarly, the asymptotic slope at *ϕ* → 1 (Eq. (E5)) reduces to

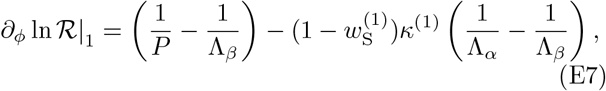

where

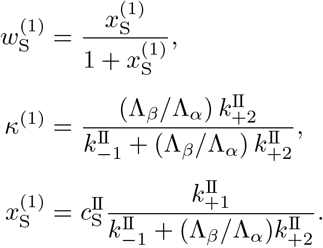

#### Onset of optimal enhancement

For the co-localization parameter regime considered in this work, ℛ exhibits either monotonic behaviour or a single maximum with respect to condensate size (Figs. 5 and 8). The corresponding optimality conditions were introduced in Eq. (34). Specifically, the large-condensate slope ∂_*ϕ*_ ln ℛ|_1_ (Eq. (E7)) remains negative over the entire range of catalytic-step enhancements explored numerically (Fig. 8). Consequently, within the co-localization regime considered here, the onset of optimal enhancement is controlled entirely by the sign of the small-condensate slope ∂_*ϕ*_ ln ℛ|_0_ (Eq. (E6)). The transition between monotonic and non-monotonic regulation occurs when the small-condensate slope changes sign, ∂_*ϕ*_ ln ℛ|_0_ = 0, which yields the critical catalytic-step kinetic partition coefficient 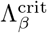 given in Eq. (35a).

When substrate unbinding dominates over catalytic turnover 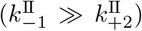, the catalytic commitment term *κ*^(0)^ becomes small. Consequently, terms proportional to *κ*^(0)^ become negligible, such that Eq. (E6) reduces to ∂_*ϕ*_ ln ℛ|_0_ ≃ Λ_*β*_ − *P* . Accordingly, in the weak catalytic-commitment regime, the critical value 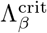 reduces to the simple criterion 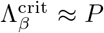 (Eq. (35b)). Notably, the kinetic parameters considered throughout this work satisfy 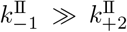 (and therefore *κ*^(0)^ ≪ 1), placing the system in this regime (Table I).

#### Interpretation of 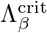

For 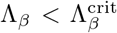, the effects of concentration partitioning dominate over the kinetic enhancement of the catalytic step. Consequently, the first-order correction to the homogeneous limit is negative and the relative initial rate ℛ decreases upon formation of a small condensate. By contrast, for 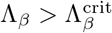, the kinetic enhancement of the catalytic step dominates over the effects of concentration partitioning, the first-order correction becomes positive, and ℛ increases upon formation of a small condensate. This transition is illustrated in Fig. 8, where ∂_*ϕ*_ ln ℛ|_0_ changes sign at 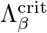.

### d. Enzyme-substrate spatial segregation regime

The richer behaviour of the segregation regime originates from the fact that both asymptotic slopes can change sign as a function of Λ_*β*_. By contrast, in the co-localization regime only the small-condensate slope changes sign, while the large-condensate slope remains negative. The emergence of additional sign combinations between the two asymptotic limits therefore generates the four distinct regulatory regimes observed in Fig. 6.

## Appendix F Parameters

## Notes

### Competing Interest Statement

The authors have declared no competing interest.

